# Development and characterization of new patient-derived xenograft (PDX) models of osteosarcoma with distinct metastatic capacities

**DOI:** 10.1101/2023.01.19.524562

**Authors:** Courtney R. Schott, Amanda L. Koehne, Leanne C. Sayles, Elizabeth P. Young, Cuyler Luck, Katharine Yu, Alex G. Lee, Marcus R. Breese, Stanley G. Leung, Hang Xu, Avanthi Tayi Shah, Heng-Yi Liu, Aviv Spillinger, Inge H. Behroozfard, Kieren D. Marini, Phuong T. Dinh, María V. Pons Ventura, Emma N. Vanderboon, Florette K. Hazard, Soo-Jin Cho, Raffi S. Avedian, David G. Mohler, Melissa Zimel, Rosanna Wustrack, Christina Curtis, Marina Sirota, E. Alejandro Sweet-Cordero

**Affiliations:** Division of Oncology, Department of Pediatrics, University of California, San Francisco, CA, USA; Department of Pathobiology, Ontario Veterinary College, University of Guelph, Guelph, ON, Canada; Bakar Computational Health Sciences Institute, University of California San Francisco. San Francisco, CA, USA; Departments of Genetics and Medicine, Stanford University School of Medicine, Stanford University, Stanford, CA, USA; Department of Pathology, Stanford University School of Medicine, Stanford University, Stanford, CA, USA; Department of Pathology, University of California San Francisco, San Francisco, CA, USA; Department of Orthopedic Surgery, Stanford University School of Medicine, Stanford University, Stanford, CA, USA; Department of Orthopedic Surgery, University of California San Francisco, San Francisco, CA, USA

## Abstract

Models to study metastatic disease in rare cancers are needed to advance preclinical therapeutics and to gain insight into disease biology, especially for highly aggressive cancers with a propensity for metastatic spread. Osteosarcoma is a rare cancer with a complex genomic landscape in which outcomes for patients with metastatic disease are poor. As osteosarcoma genomes are highly heterogeneous, a large panel of models is needed to fully elucidate key aspects of disease biology and to recapitulate clinically-relevant phenotypes. We describe the development and characterization of osteosarcoma patient-derived xenografts (PDXs) and a panel of PDX-derived cell lines. Matched patient samples, PDXs, and PDX-derived cell lines were comprehensively evaluated using whole genome sequencing and RNA sequencing. PDXs and PDX-derived cell lines largely maintained the expression profiles of the patient from which they were derived despite the emergence of whole-genome duplication (WGD) in a subset of cell lines. These cell line models were heterogeneous in their metastatic capacity and their tissue tropism as observed in both intravenous and orthotopic models. As proof-of-concept study, we used one of these models to test the preclinical effectiveness of a CDK inhibitor on the growth of metastatic tumors in an orthotopic amputation model. Single-agent dinaciclib was effective at dramatically reducing the metastatic burden in this model.

**Graphical Abstract:** 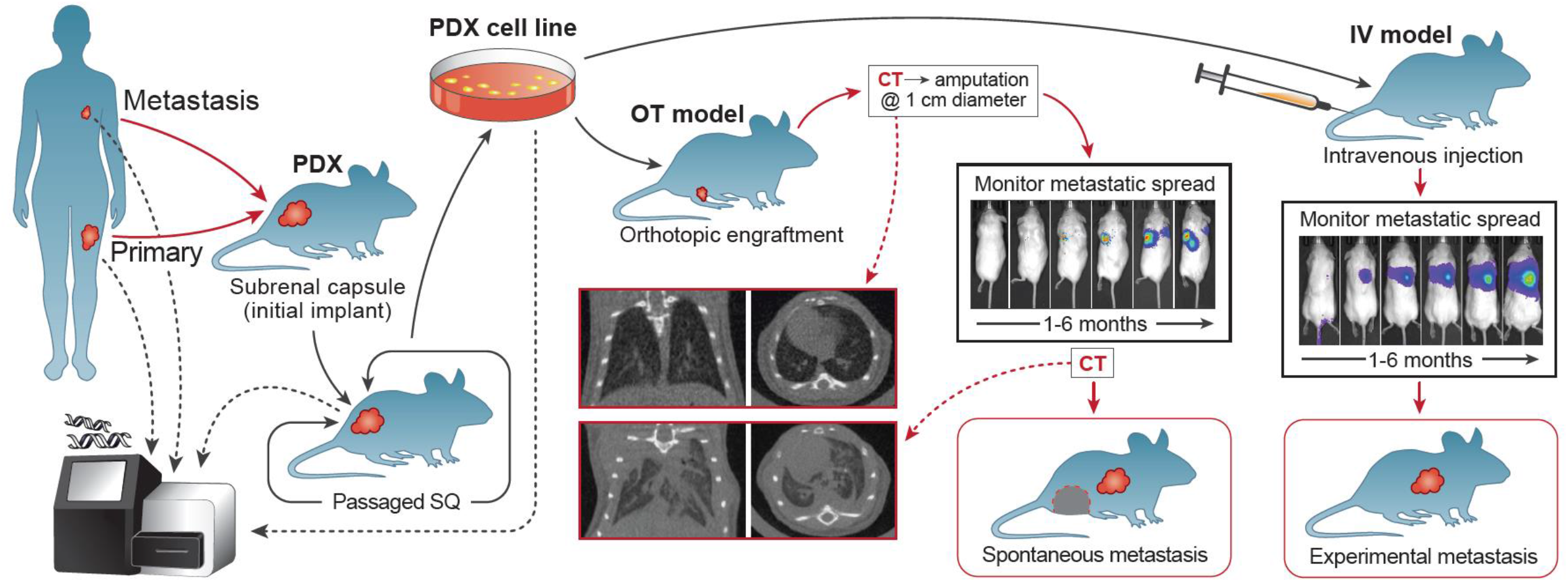

## Introduction

Osteosarcoma is a rare malignant bone tumor most commonly diagnosed in children, adolescents, and young adults. The majority of patients have micrometastases at diagnosis, requiring the use of intensive adjuvant chemotherapy (1–3). Despite aggressive treatment, progression often presents as distant metastatic disease in the lungs or bone, and patients who develop metastases have a 5-year survival rate of 20-30% (3–7). Over the past several decades, there has been limited progress in identifying new therapies for patients with osteosarcoma, and survival has not substantively improved, particularly for patients with metastatic disease (7–9). This lack of progress may be partly due to a paucity of models that accurately recapitulate osteosarcoma metastatic progression. Osteosarcoma genomes are characterized by multiple copy number alterations, chromothripsis, and aneuploidy, resulting in significant heterogeneity in oncogenic drivers. Well-characterized models are crucial not only for elucidating mechanisms of metastasis but also for identification of new therapeutic strategies (10, 11).

While *in vitro* assays are often used to predict or inform on the metastatic capacity of neoplastic cells (e.g. invasion, migration, anoikis, etc.), *in vivo* modeling remains the gold standard to fully evaluate the metastatic cascade. Metastasis models where cancer cells are disseminated after intravenous (IV) injection recapitulate the latter half of the metastatic cascade, including survival within the bloodstream, extravasation, tissue colonization, and tumor establishment. However, to model the full metastatic cascade, models that metastasize spontaneously from a relevant orthotopic (OT) site are required.

Only a limited number of established osteosarcoma cell lines have demonstrated the ability to consistently form metastasis when injected IV (12–20) (Supplemental Table S1). To develop additional models, prior work has sought to increase metastatic capacity by exposure to carcinogens, introduction of oncogenes, or serial passaging of established osteosarcoma cell lines through mice (14, 15, 21–27). These strategies have successfully enhanced the metastatic capacity of several osteosarcoma cell lines in both IV and OT models (12, 14). However, these approaches may introduce alterations that are not reflected in endogenous human metastasis. Furthermore, murine OT osteosarcoma models are most commonly generated by implanting tumor cells/tissue into the epiphyseal bone or onto the periosteal surface of the tibia (12, 16, 19, 20, 27–41). However, injecting tumor cells/tissue directly into the bone can lead to immediate exit through the vasculature, essentially skipping the initial stages of the metastatic cascade and seeding the lungs directly (42–45). The periosteal implantation strategy to model spontaneous metastasis requires tumor cells to invade through the cortex without risk of inadvertently introducing cells into the vasculature at implantation. Models incorporating amputation of the affected limb, provide additional time for the development of metastasis in the lungs or other organs (12, 16, 36, 38, 41). Since surgical removal of the primary tumor is a component of the standard of care for patients with osteosarcoma, inclusion of primary tumor removal in a mouse model more closely recapitulates management of the human disease.

The use of patient-derived xenografts (PDXs) to model cancer enhances the probability of retaining key features of the human disease (11, 42, 46–49). However, only a limited number of osteosarcoma PDX models have been evaluated for metastatic potential *in vivo* (30, 36, 40, 42, 50–52). Cell lines generated from PDXs may also be useful to establish more consistent and more tractable models for the exploration of molecular pathogenesis and for testing novel therapeutic drugs for pediatric osteosarcoma compared to established commercially available cell lines, many of which have been in culture for decades (53).

To enable the development of new models to study osteosarcoma metastasis, we generated PDXs from clinically annotated patient tumors. Cell lines were then generated from the PDXs and rigorously characterized *in vivo* using both an intravenous and an orthotopic model (see Graphical Abstract). Whole genome sequencing (WGS) and RNA sequencing (RNAseq) were used to evaluate consistency between the PDXs and the patient tumor from which they were derived. Here, we describe the *in vivo* features of these novel PDX-derived cell lines including tumorigenicity and metastatic capacity in both tail vein IV and paratibial OT amputation models. Bioluminescence imaging (BLI) was used to quantify whole body metastatic burden, while immunohistochemistry (IHC) was used to aid quantification of metastatic tumor burden in the lung and liver. Furthermore, to demonstrate the utility of PDX-derived cell lines for preclinical testing, we performed a proof-of-concept study which showed that a CDK inhibitor decreased metastatic tumor growth of a MYC amplified cell line after amputation of an OT xenograft. We expect that these PDX-derived cell line models will be broadly useful to researchers investigating mechanisms of osteosarcoma metastasis as well as the preclinical development of therapies to treat metastatic disease.

## Results

### Osteosarcoma PDXs are highly similar to their patient of origin

Tumor samples obtained from diagnostic biopsies, post-chemotherapy primary resections, or metastases, were collected directly from patients at the time of surgery with institutional review board approval and appropriate consent. Tumor fragments were implanted into the subrenal capsule of immunocompromised mice to generate PDXs. PDXs retain the morphological features of their respective patient tumor based on histology (11), even over multiple passages (Supplemental Figure S1). From these PDXs, we generated eleven PDX-derived cell lines. Of these cell lines, three were obtained from pre-treatment biopsies, three from post-treatment resections, and four from distant metastasis (Table 1).

**Table 1:** Osteosarcoma PDX-derived cell line patient origin details, genomic features, and *in vivo* characteristics

| ID | Patient |  |  |  | Cell Line |  |  |  | Met *in vivo* |  | Xenograft |  |
| --- | --- | --- | --- | --- | --- | --- | --- | --- | --- | --- | --- | --- |
|  | Age (yr) | Sex | Met @ Dx | Met Development | Origin | Tx Status | Site Origin | Genomic Feature | IV | OT | Take | Days to Amp |
| OS052 | 17 | F | Yes | N/A | primary | treated | fibula | PTEN loss | 100% | 22% | 100% | 35 |
| OS152 | 11 | F | No | Yes | metastatic | treated | scapula | MYC amplification | 100% | 100% | 100% | 39 |
| OS186 | 13 | M | Suspected | Yes | metastatic | treated | mandible | MYC & CCNE1 amplification | 0% | 0% | 100% | 83 |
| OS384 | 12 | F | No | No @ 71 months | primary | treated | tibia | VEGFA amplification | 100% | 88% | 100% | 119 |
| OS457 | 17 | M | Yes | N/A | primary | naïve | humerus | CCNE1 amplification | 0% | N/A | 0% | N/A (>168) |
| OS525 | 20 | F | Yes | N/A | metastatic | treated | lung | AKT1 amplification | 100% | 100% | 100% | 53 |
| OS526 | 13 | M | Yes | N/A | primary | naïve | tibia | CCNE1 amplification | 100% | 0% | 100% | 144 |
| OS742 | 8 | F | No | Yes | primary | naïve | tibia | MYC & CDK4 amplification | 100% | 100% | 100% | 30 |
| OS766 | 30 | F | Suspected | Yes | metastatic | treated | femur | CDK4 amplification | TBD | TBD | TBD | TBD |
| OS833 | 14 | M | No | Yes | primary | treated | fibula | MYC amplification | TBD | TBD | TBD | TBD |

Amp = amputation, Dx = diagnosis, IV = intravenous, Met = metastasis, N/A = not applicable, OT = orthotopic, TBD = to be determined, Tx = treatment

Where feasible, fresh-frozen tumor samples were analyzed by WGS and RNAseq. WGS and RNAseq was also performed for all PDXs and PDX-derived cell lines (Figures 1–2).

**Figure 1.**
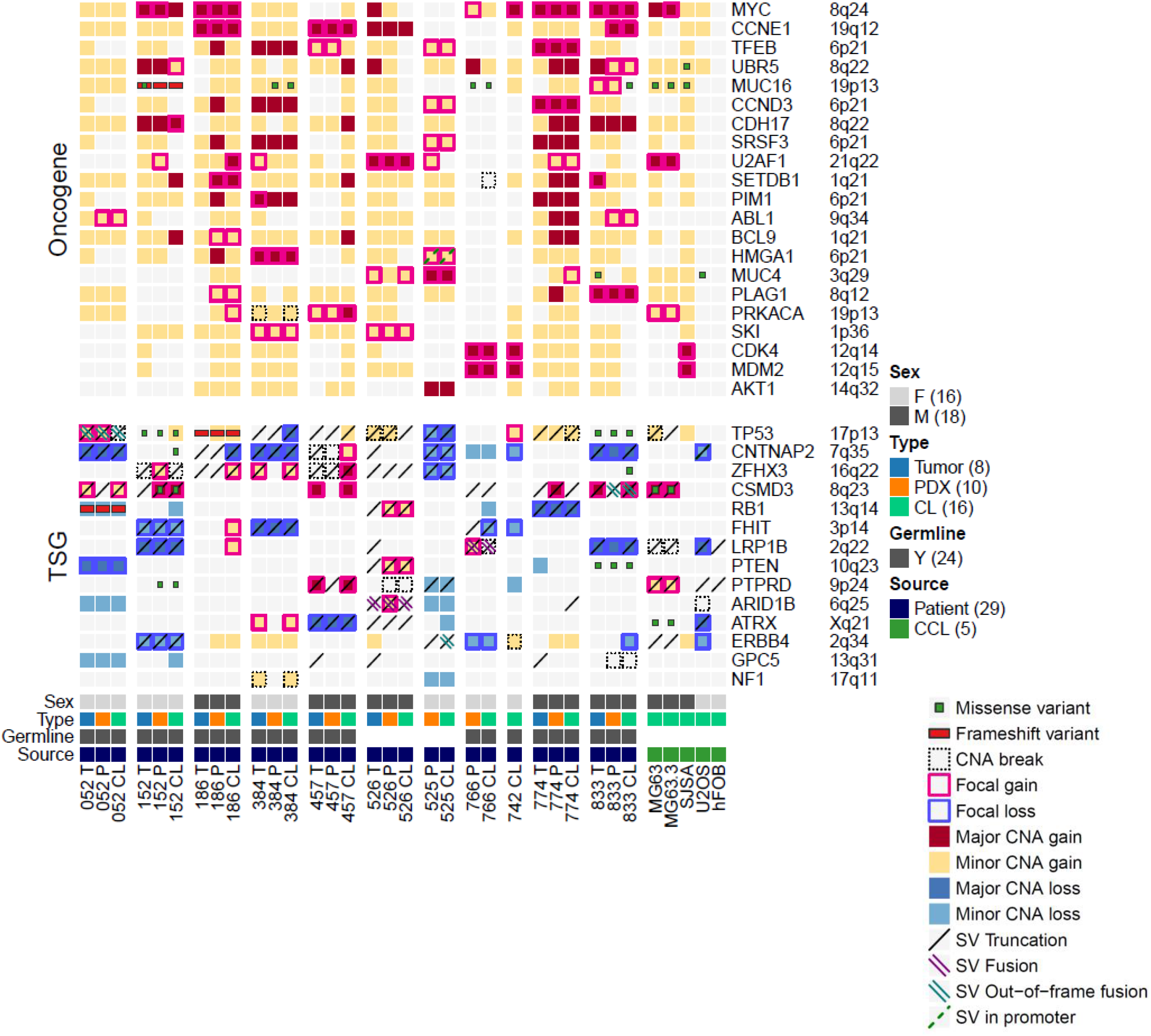
Whole genome sequencing of patient, patient-derived xenograft (PDX), and PDX-derived cell line. Oncoprint showing the genomic alterations in oncogenes and tumor suppressor genes (TSGs) across primary tumors (T), PDX (P), PDX-derived cell lines (CL), and commercially available cell lines (CCL). Alterations in top recurrently altered and certain selected oncogenes and tumor suppressors are shown. Focal copy-number alterations are defined as contiguous segments of similar copy-number change less than 1Mb in length. SV fusion calling and frame-ness determined using WGS only without RNAseq confirmation.

**Figure 2.**
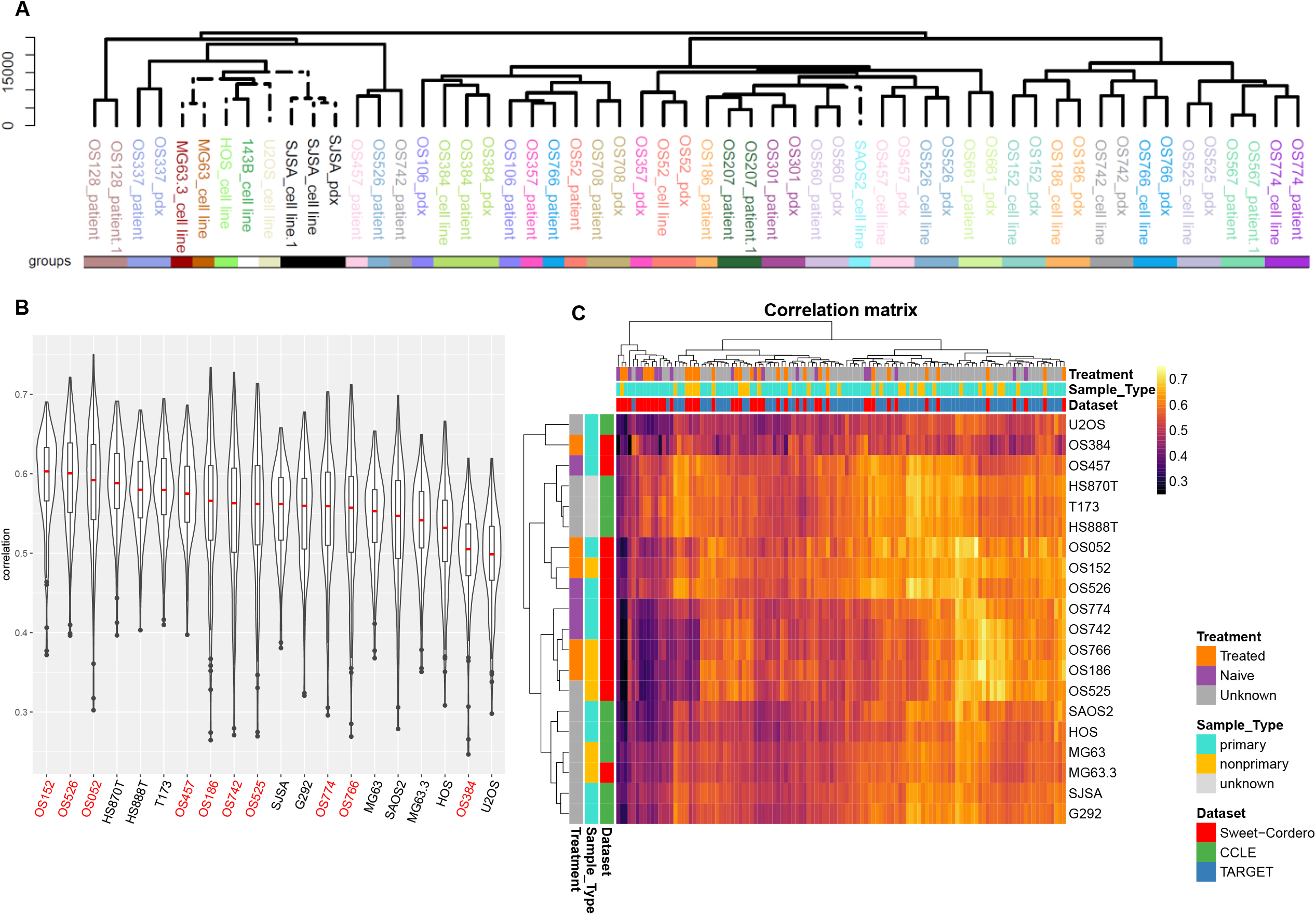
Sequencing of osteosarcoma patients, patient-derived xenografts (PDXs), and PDX-derived cell lines. **(A)** Patient tumors and their corresponding PDX and PDX-derived cell line exhibit remarkable similarity and cluster separately to established commercially available cell lines. Hierarchical clustering of RNAseq data from osteosarcoma patient tumors, PDXs, PDX-derived cell lines, and established commercially available cell lines. **(B)** Gene expression of osteosarcoma PDX-derived cell lines correlates with osteosarcoma patient tumors. Violin plot of Spearman’s correlations between all possible pairs of primary patient tumor samples (CCLE and Sweet-Cordero lab) and either established commercially available osteosarcoma cell lines (CCLE and Sweet-Cordero lab) or PDX-derived cell lines (Sweet-Cordero lab) using the 5000 most variable genes, separated by individual cell line (*x*-axis). Names of cell lines from the Sweet-Cordero lab are in red and names of cell lines from the CCLE are in black. The median correlation coefficients are depicted by the red center line and the overlaid boxplots depict the upper and lower quartiles, with the whiskers depicting 1.5 times the IQR. **(C)** Heatmap showing the Spearman’s correlations between osteosarcoma cell lines (rows) and primary tumor samples (columns). The color annotation bars on the x- and y-axis indicate the treatment status, tumor type (e.g. primary, nonprimary, unknown), and dataset of each sample.

Unsupervised hierarchical clustering of RNAseq data demonstrated a remarkable similarity of the patient tumor to the corresponding PDX and PDX-derived cell line (Figure 2A). In contrast, most established commercially available osteosarcoma cell lines (HOS, 143B, SJSA, U2OS, MG63, and MG63.3) clustered separately from our cohort of patient samples, PDXs, and PDX-derived cell lines (Figure 2A). Thus over decades of *in vitro* passaging, these established commercially available cell lines may have drifted away from patient samples, providing support for the hypothesis that newly derived PDX-derived cell lines retain key features of the gene expression programs of primary tumors. Next, we compared the gene expression profiles of PDX-derived cell lines and established commercially available cell lines to patient primary tumor samples from publicly available data (Figure 2B-C). Overall, the PDX-derived cell lines had a higher correlation to human osteosarcoma patient samples. For example, across all cell lines tested, the three with the highest mean correlation to human osteosarcoma patient samples were PDX-derived cell lines, whereas the mostly commonly used commercial osteosarcoma cell lines are the least correlated (Figure 2B).

Osteosarcoma is genomically complex, with widespread aneuploidy and structural rearrangements being hallmarks for this cancer (7, 54). We previously described comparison of a limited panel of osteosarcoma PDXs to their corresponding tumor of origin (11). Here we sought to characterize the changes that occurred at the genomic level when comparing this new collection of PDX-derived cell lines to their matched patient and PDX samples. We compared the copy number profile of patient tumors and their corresponding PDXs and PDX-derived cell lines. Overall, we found a high degree of concordance of copy number profiles as identified by WGS (Supplemental Figure S2). In two of the cell lines tested *in vivo* (OS457 and OS186), whole-genome duplication (WGD) led to a wider degree of difference compared to the tumor of origin as quantified by the percentage of genome segments overlapping with regards to copy number (Supplemental Figure S2A). In one case (OS457), the WGD appears to have occurred in the establishment of the PDX (Supplemental Figure S2A). In the second case (OS186), the WGD occurred sometime after the establishment of the cell line (Supplemental Figure S2A). Similarly, Circos plots demonstrated high similarity in the structural rearrangements between patient, PDX, and PDX-derived cell line (Supplemental Figure S3).

### Metastatic capacity of osteosarcoma PDXs is highly variable

Next, we evaluated the metastatic capacity of these PDXs *in vivo*. Differences in metastatic potential was assessed using several different methods. Lungs from a subset of mice harboring subcutaneous (SQ) xenografts, during PDX generation and passaging, were evaluated for the presence of spontaneous metastasis. We also characterized the ability of a subset of these PDXs to form metastases after IV injection. Lungs were collected at the time of PDX passaging for mice with SQ PDX implants and 28 days post-injection for mice injected with PDX cells IV. Evidence of metastasis was evaluated histologically revealing the presence of both micro- and macro-metastases (Supplemental Figure S4A-C). Of the PDXs evaluated for spontaneous metastasis from SQ tumors, eight of eleven demonstrated the ability to form lung metastasis (Supplemental Figure S4B). When injected IV, three of five PDXs induced lung metastasis (Supplemental Figure S4C). We observed some discordance in the ability of certain PDXs to form metastasis during SQ passaging versus in the IV model. For example, OS128 induced metastasis in the IV model but not during SQ passaging, while OS152 and OS186 metastasized during SQ passaging, but not after IV administration (Supplemental Figure S4B-C).

### Osteosarcoma PDX-derived cell lines form metastases when injected IV

As we noted a high degree of correlation at the level of transcription between the PDX and the PDX-derived cell lines (Figure 2A), we reasoned that the later would be a more tractable model system for metastatic studies. To assess the metastatic capacity of osteosarcoma PDX-derived cell lines *in vivo*, PDX-derived cell lines were transduced with a lentiviral vector expressing luciferase to facilitate dynamic monitoring of metastatic burden in NSG mice. We first determined that proliferation rates between luciferase-expressing PDX-derived cell lines and the parental version were similar (Supplemental Figure S5A). Next, cells were introduced via the lateral tail vein and mice were monitored with BLI to estimate whole body metastatic burden (Figure 3A). Most (6/8) osteosarcoma PDX-derived cell lines tested generated metastases with increasing BLI signal over time (Figure 3B-C and Supplemental Figure S6A-H). Study duration for each cell line ranged from one to a maximum of six months and exhibited a trend of being inversely correlated with BLI signal at endpoint (Table 2 and Figure 3B-C). A subset of the parental (non-luciferase-expressing) PDX-derived cell lines were also evaluated using the IV model (Supplemental Figure S5B-E and Supplemental Table S2).

**Figure 3.**
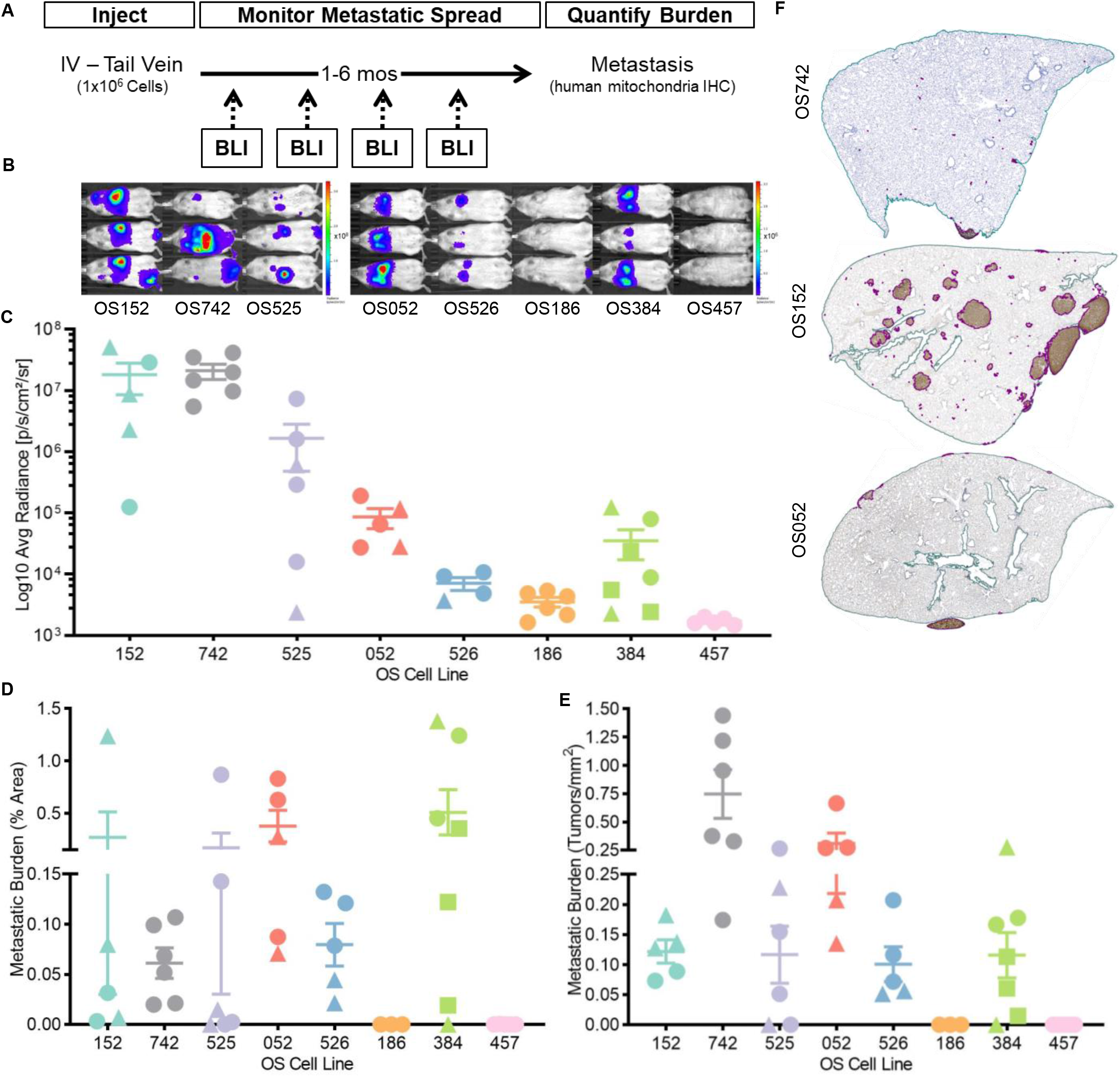
Osteosarcoma patient-derived xenograft (PDX)-derived cell lines metastasize after intravenous (IV) injection. IV metastatic capacity was determined by injecting 1 x 10^6^ cells into the lateral tail vein of 2-5-month-old NSG mice. For all graphs, within each cell line, mice implanted at the same time are represented by the same symbol. **(A)** IV model schematic: after IV injection, mouse health was monitored for clinical signs associated with metastasis and bioluminescence imaging (BLI) was performed monthly to estimate burden to determine study endpoint for each cell line. At endpoint, tissues were harvested for quantitative burden analysis. **(B)** Metastatic burden varied between cell lines. Representative images from 3 mice per cell line of the BLI signal at endpoint representing total tumor burden from all sites. **(C)** Total BLI signal at endpoint, ordered from fastest to slowest time to endpoint, for all mice for each PDX-derived cell line. **(D-E)** Cell lines most frequently metastasized to lung. Metastatic burden quantification of lung tissue immunolabeled with human mitochondria antibody for overall tumor area (D) and number of tumors per mm^2^ (E) for individual mice. **(F)** Tumor size and distribution in the lung parenchyma varied between cell lines. Microscopic images of lung immunolabeled with human mitochondria antibody demonstrating metastatic burden.

**Table 2:** Osteosarcoma PDX-derived cell line *in vivo* intravenous (IV) study summary table

|  |  | OS152 | OS742 | OS525 | OS052 | OS526 | OS186 | OS384 | OS457 |
| --- | --- | --- | --- | --- | --- | --- | --- | --- | --- |
| INTRAVENOUS |  |  |  |  |  |  |  |  |  |
| # Mice Total [# Studies] |  | 5 [2] | 6 [1] | 6 [2] | 5 [2] | 5 [2] | 6 [2] | 6 [3] | 5 [1] |
| Endpoint (avg days) |  | 53 | 55 | 80 | 85 | 140 | 160 | 168 | 168 |
| Endpoint Decision with CS, if Applicable |  | BLI, CS | BLI, CS | CS | BLI | BLI | max | max | max |
|  |  | abd mass | low BCS, gait | paralysis | N/A | N/A | N/A | N/A | N/A |
| Metastatic Sites (%) | Lung | 100 | 100 | 67 | 100 | 100 | N/A | 100 | N/A |
|  | Bone | 20 | 33 | 100 | 20 | 0 |  | 0 |  |
|  | Liver | 60 | 100 | 67 | 0 | 0 |  | 0 |  |
|  | Other | 20 | 33 | 33 | 20 | 0 |  | 0 |  |

abd = abdominal, avg = average, BCS = body condition score, BLI = bioluminescence imaging, CS = clinical signs

A wide range of tropism with regards metastatic sites was observed (Table 2). All 6 metastatic PDX-derived cell lines induced pulmonary metastases: 5/6 metastasized consistently to lung in every mouse and 1/6 did so frequently (67%; Table 2 and Figure 3D-E). Lung metastases tended to be higher in occurrence and larger in size toward the pleural surface (Figure 3F). Four cell lines metastasized to bone (spine, sternum, rib, skull, and long bone) in 20-100% of mice per PDX-derived cell line. Lung and bone are the two most common sites of metastasis in osteosarcoma patients. Liver metastasis (which is uncommon in patients) was also observed for three PDX-derived cell lines in the IV model (Supplemental Figure S7A-C), although never in isolation. The non-luciferase versions tested, metastasized to the same sites as BLI enabled lines with minor differences in location and burden (Supplemental Figure S5B-E and Supplemental Table S2). The three most aggressively metastatic cell lines (OS152, OS742, and OS525) formed metastases in all mice tested, resulting in the highest overall burden, greatest number of sites, and shortest latency compared to the remaining PDX-derived cell lines (Table 2, Figure 3, and Supplemental Figures S6-7). With most PDX-derived cell lines demonstrating the ability to metastasize in an IV model with high incidence but low burden, we next tested their capacity to spontaneously metastasize in an orthotopic (OT) amputation model.

### Osteosarcoma PDX-derived cell lines are tumorigenic in an OT model

We sought to develop an OT osteosarcoma model to fully evaluate the entire metastatic cascade as a clinically relevant model for preclinical testing. Paratibial implantation of luciferase-expressing PDX-derived cells into NSG mice was performed to assess tumorigenicity and capacity for spontaneous metastasis (Figure 4A). All but one PDX-derived cell line (OS457) formed xenografts within six months of implantation (Figure 4B and Supplemental Figure S8A-H). This cell line was the most divergent from both the patient tumor and PDX from which it was derived, likely due to a WGD event that occurred while the cell line was established from the PDX (Supplemental Figure S2A). Time to amputation (when the xenograft reached 1 cm diameter), ranged from less than one month to eight months for individual mice whereas average time to amputation for each cell line ranged from one to five months (Table 3, Figure 4B, and Supplemental Figure S8A-G). Each tumorigenic PDX-derived cell line demonstrated a consistent growth rate for OT xenografts except OS384 which varied widely (Figure 4B and Supplemental Figure S8G). For another PDX-derived cell line (OS457), BLI signal was detected after six months at the OT implant site for all mice (Supplemental Figure S8I), but no xenografts could be palpated nor were they captured histologically. PDX-derived cell line xenografts exhibited heterogeneous histologic phenotypes (Supplemental Figures S8J), similar to what is seen in human patients. Most PDX-derived cell line xenografts exhibited regions with similar histologic phenotypes to the patient tumors from which they were derived (Supplemental Figures S8J).

**Figure 4.**
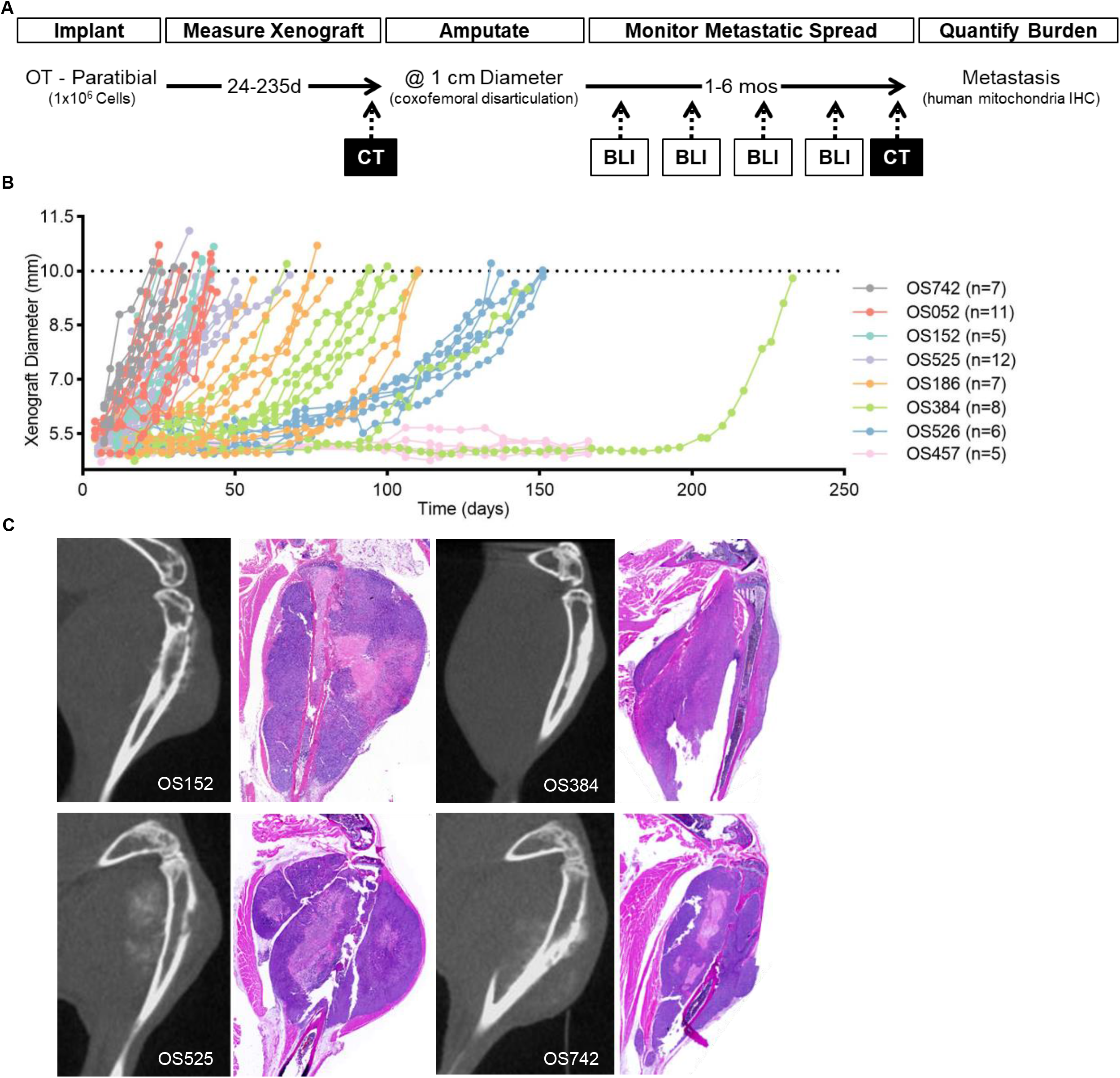
Osteosarcoma patient-derived xenograft (PDX)-derived cell lines form invasive xenografts after paratibial orthotopic (OT) implantation. **(A)** OT model schematic: 1 x 10^6^ cells were implanted paratibially into the right hindlimb of 2–5-month-old NSG mice. Limb diameter was measured bi-weekly until reaching 1 cm when limb microCT was performed, followed by amputation via coxo-femoral disarticulation. After amputation, mouse health was monitored for clinical signs associated with metastasis and bioluminescence imaging (BLI) was performed monthly to estimate burden and to determine study endpoint for each cell line. At endpoint, tissues were harvested for quantitative burden analysis. **(B)** Mice from most cell lines required amputation at similar time points. Bi-weekly xenograft diameter measurements for individual mice implanted with each PDX-derived cell line. All but one cell line, OS457, formed xenografts that reached 1 cm in diameter. **(C)** After paratibial implantation, cell line xenografts invaded through the cortex. Example hindlimb microCTs with corresponding histopathology demonstrating medullary cavity and trabecular bone invasion and tumor expansion for several tumorigenic PDX-derived cell lines. Hematoxylin and eosin.

**Table 3:** Osteosarcoma PDX-derived cell line *in vivo* orthotopic (OT) study summary table

|  |  | OS742 | OS152 | OS525 | OS052 | OS186 | OS384 | OS526 | OS457 |
| --- | --- | --- | --- | --- | --- | --- | --- | --- | --- |
| ORTHOTOPIC |  |  |  |  |  |  |  |  |  |
| # Mice Total [# Studies] |  | 5 [1] | 7 [1] | 6 [2] | 9 [3] | 8 [2] | 5 [2] | 7 [2] | 5 [1] |
| Amp Time (avg days) |  | 39 | 30 | 53 | 35 | 119 | 144 | 83 | N/A |
| Endpoint (avg days) |  | 29 | 32 | 84 | 168 | 168 | 161 | 168 | 168 |
| Endpoint Decision with CS, if Applicable |  | BLI | BLI, CS | BLI, CS | max | max | max, CS | max | N/A |
|  |  | N/A | abd mass | paralysis | N/A | N/A | low BCS | N/A | N/A |
| Metastatic Sites (%) | Lung | 100 | 100 | 50 | 0 | 88 | N/A | N/A | N/A |
|  | Bone | 40 | 43 | 100 | 11 | 0 |  |  |  |
|  | Liver | 100 | 100 | 50 | 0 | 0 |  |  |  |
|  | Other | 0 | 86 | 50 | 22 | 13 |  |  |  |

abd = abdominal, amp = amputation, avg = average, BCS = body condition score, BLI = bioluminescence imaging, CS = clinical signs

Cell line xenograft growth was similar regardless of the presence of luciferase, with a tendency for the luciferase-expressing version to reach amputation more slowly after implantation (Supplemental Figure S9A-D). microCT prior to amputation detected tibial osteolysis and medullary cavity or trabecular bone invasion for all tumorigenic PDX-derived cell line xenografts in the OT model, which was confirmed histologically (Figure 4C), highlighting the aggressiveness of these models.

### Osteosarcoma PDX-derived cell lines spontaneously metastasize in an OT amputation model

Post-amputation, mice in OT studies were monitored for metastatic burden for up to six months using BLI and health monitoring (Table 3, Figure 4A, Figure 5A-B and Supplemental Figure S10A-G). Study duration after amputation ranged from one to a maximum of six months (Table 3 and Supplemental Figure S10A-G). Spontaneous metastases developed in mice for five of the seven xenograft forming PDX-derived cell lines with BLI signal increasing over time (Table 3 and Supplemental Figure S10A-G). BLI signal at endpoint showed a trend of being inversely correlated with time to endpoint (Table 3 and Figure 5A-B) demonstrating that higher metastatic burden causes increased morbidity *in vivo*.

**Figure 5.**
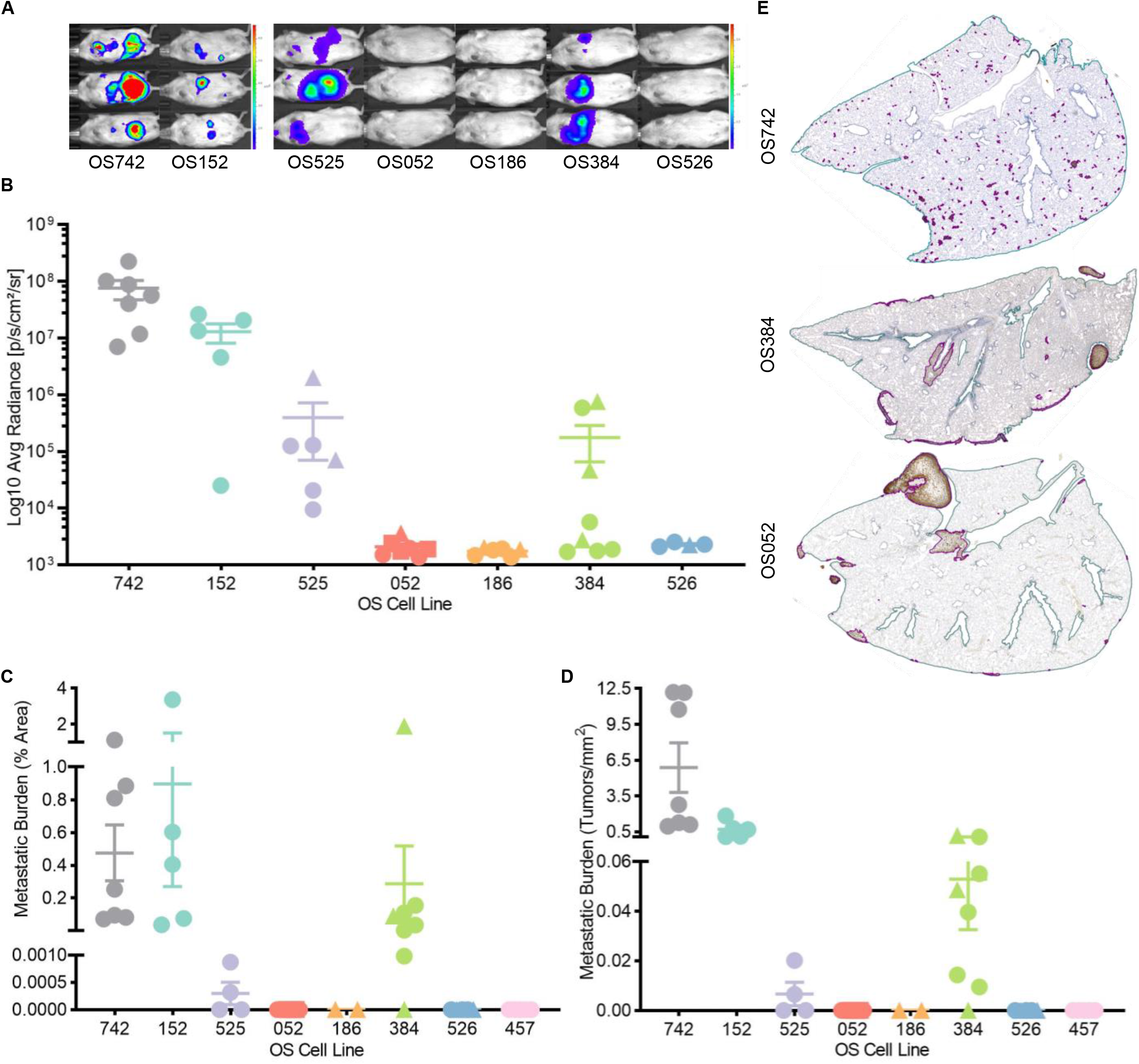
Osteosarcoma patient-derived xenograft (PDX)-derived cell lines metastasize spontaneously in an orthotopic (OT) amputation model. For all graphs, within each cell line, mice implanted at the same time are represented by the same symbol. **(A)** Metastatic burden varied between cell lines. Representative images of the BLI signal at endpoint for each cell line representing total tumor burden from all sites. **(B)** Total BLI signal at endpoint for all mice, ordered from fastest to slowest time to amputation, for each PDX-derived cell line. **(C-D)** Metastases were observed most consistently in the lung. Metastatic burden quantification of lung tissue immunolabeled with human mitochondria antibody for overall tumor area (C) and number of tumors per mm^2^ (D) for individual mice. **(E)** Tumor size and distribution in lung parenchyma varied between cell lines. Microscopic images of lung immunolabeled with human mitochondria antibody demonstrating metastatic burden.

As with the IV model described above, there was a high degree of tissue-specific tropism in metastasis development (Table 3 and Figure 5A). Two of the five metastatic PDX-derived cell lines (OS742 and OS152) metastasized consistently to the lung in all mice (Table 3 and Figure 5C-D). Of the remaining three xenograft-forming cell lines that metastasized, two (OS525 and OS384) metastasized to lung with high frequency (50 and 88%, respectively) and one (OS052) never metastasized to the lung (Table 3 and Figure 5C-D). PDX-derived cell lines OS152, OS525, and OS742 also metastasized to bone (spine, skull, and long bone) as well as liver in up to 100% of mice (Table 3 and Supplemental Figures S10H-I and S11). Four of the PDX-derived cell lines also metastasized to other soft tissue sites (abdomen, kidney, axillary region, and mediastinum) in 13-86% of mice (Table 3). Lung burden by area for all mice remained below 4%, whereas liver burden reached as high as nearly 50% (Figure 5C and Supplemental Figure S11A). The three most aggressive cell lines (OS152, OS742, and OS525) metastasized in all mice with the highest overall burden, to the most sites and with the shortest latency (Table 3, Figure 5, and Supplemental Figures S10-11) compared to the remaining PDX-derived cell lines. OS052 was one of the fastest cell lines to grow a 1 cm xenograft at the OT site but had low capacity to form spontaneous metastasis (Table 3, Figures 4B and 5A-B). Using the OT model, metastatic capacity and site tropism between the luciferase and parental versions were similar (Supplemental Figure S9E-H and Supplemental Table S2).

### Performance of established commercial osteosarcoma cell lines using IV and OT *in vivo* models

Next, we compared the *in vivo* metastatic phenotypes of the panel of osteosarcoma PDX-derived cell lines to a small subset of commercially available established osteosarcoma cell lines by evaluating them using the same IV and OT models described above.

In the IV model, SJSA and MG63.3 caused metastasis-associated morbidity faster than the most aggressive PDX-derived cell lines; the BLI signal for SJSA at endpoint was comparable to the most aggressive PDX-derived cell lines but in half of the time (Supplemental Table S4 and Supplemental Figure S12A-B). The metastatic lung burden for SJSA and MG63.3 was much higher than the PDX-derived cell lines, whereas for MG63 it was lower than many of the PDX-cell derived cell lines (Supplemental Figure S12C-D). Unlike with the PDX-derived cell lines, lung metastases formed by established commercially available osteosarcoma cell lines were more evenly distributed throughout the pulmonary parenchyma (Supplemental Figure S12F). However, MG63.3 tended to track along large airways and the MG63/MG63.3 tumors appear to contain a reduced number of human mitochondria compared to SJSA or PDX-derived cell line metastases (Supplemental Figure S12E and Figure 3F). None of the established commercially available cell lines metastasized to bone and only SJSA metastasized to the liver (Supplemental Figure S12F-H). Since human osteosarcoma frequently metastasizes to bone, a useful feature of the cell line panel described here is the observed bone tropism.

In the OT model, SJSA and MG63.3 xenografts grew quickly with all mice requiring amputation by one month; for MG63, xenografts were sporadically palpable in some mice, but limb diameter was never much larger than ~6 mm in diameter, even after > 250 days post-implantation (Supplemental Table S5 and Supplemental Figure S13A-B). Like the PDX-derived cell line xenografts, SJSA and MG63.3 xenografts invaded through the tibial cortex (Supplemental Figure S13C-D). For SJSA and MG63.3, study duration from amputation to endpoint (Supplemental Table S5) and metastatic lung burden (Supplemental Figure S13E-I) was similar to the most aggressive PDX-derived cell lines; neither of these cell lines metastasized to bone in the OT model. After characterizing the *in vivo* metastatic phenotypes of PDX-derived and established osteosarcoma cell lines in IV and OT models, we next set out to demonstrate their potential utility in genomically informed targeted drug studies.

### CDK inhibition with dinaciclib decreases metastasis in MYC amplified osteosarcoma *in vivo*

A primary rationale for the development of an osteosarcoma OT amputation model using PDX-derived cell lines that metastasize spontaneously is to use this model to test new therapeutic approaches that may be effective for treatment of metastatic disease. To this end, we tested the response to targeted therapy in one of the PDX-derived cell lines, OS152, which carries a MYC amplification (Table 1 and Figure 1). We have previously shown that MYC amplified SQ xenografts can be targeted with the CDK inhibitor dinaciclib (11). Dinaciclib inhibits CDK2/5/9 which results in downregulation of MYC expression (55). To determine the utility of this model as a preclinical tool to assess the efficacy of dinaciclib therapy, mice were implanted with OS152 luciferase-expressing cells OT as described above. Intraperitoneal (IP) dinaciclib administration (20 mg/kg 5 times/week) commenced 4 days post-amputation (Figure 6A). At endpoint, the BLI signal was significantly lower in dinaciclib treated mice (*p* = 0.0153, t test; Figure 6C-D) and the difference between the treatment and control groups increased with time (Figure 6B). In the treatment group, there was a significant reduction in the average metastatic tumor area (lung: *p* = 0.0315 and liver: *p* = 0.0040, both Mann-Whitney test; Figure 6E) and number of metastases per mm^2^ (lung: *p* = 0.0024, t test and liver: *p* = 0.0400, Mann-Whitney test; Figure 6F) in the lung and liver. These results suggest that the OT metastasis model developed here can be used to evaluate novel targeted therapeutic approaches to treat osteosarcoma metastasis.

**Figure 6.**
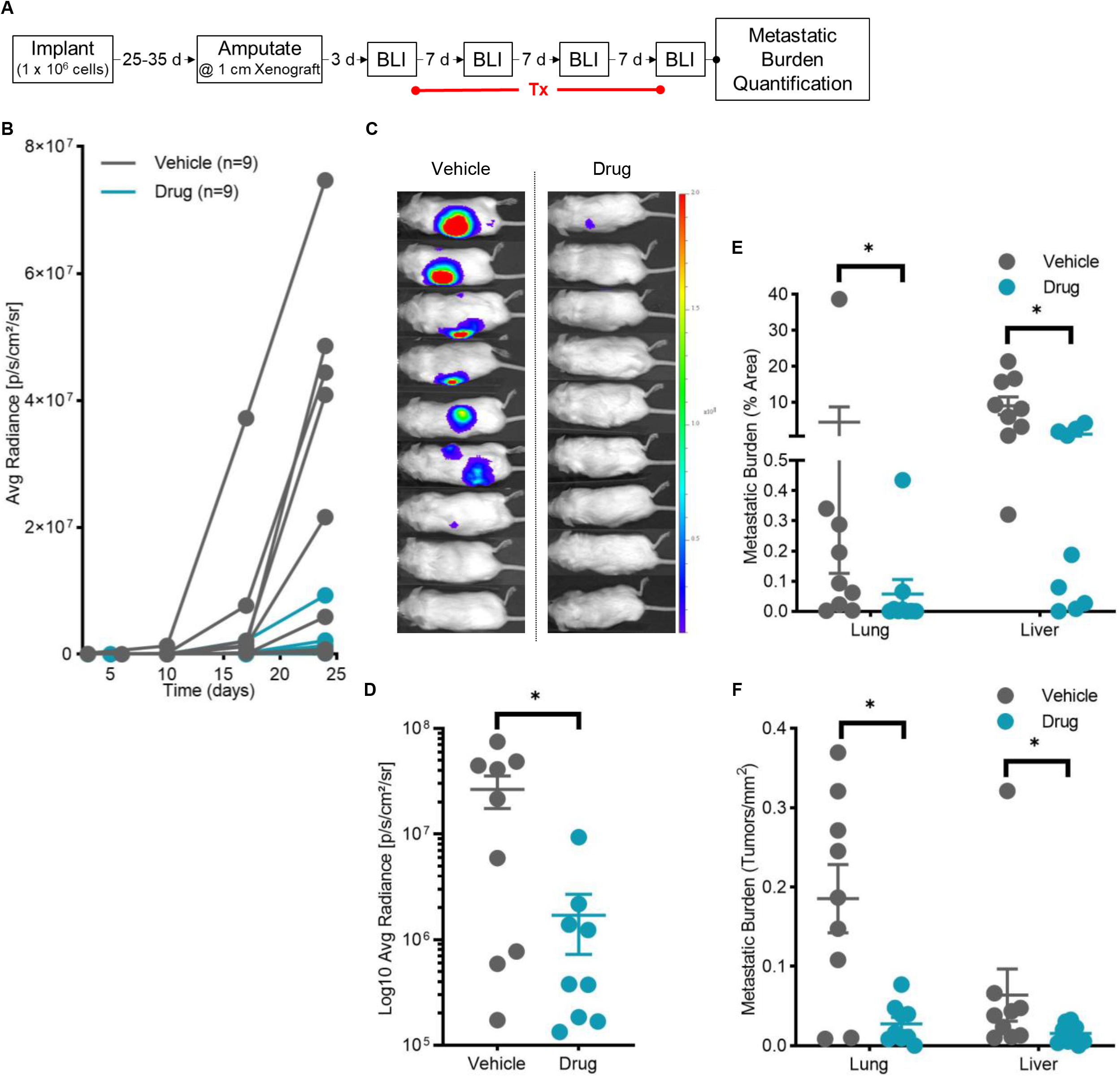
MYC amplified osteosarcoma metastases generated using an orthotopic (OT) amputation mouse model responded to targeted treatment with a CDK2/5/9 inhibitor. **(A)** Drug study schematic: OT xenografts were established by implanting 1 x 10^6^ OS152 cells paratibially into 6-9-week-old NSG mice. Xenograft diameter was measured biweekly, and the affected limb was amputated at 1 cm in diameter. Dinaciclib treatment (20 mg/kg 5 times/week IP) was initiated 3 days post-amputation and bioluminescence imaging (BLI) was performed weekly to monitor metastatic burden. At endpoint, tissues were harvested for quantitative burden analysis. **(B)** Tumor burden increased over time in both groups but was higher in the vehicle only group. Weekly BLI signal for all individual mice over the duration of the study. **(C)** At endpoint, BLI signal was higher in the vehicle only group. BLI signal images for individual mice at endpoint. **(D)** Total BLI signal at endpoint for all mice representing whole body metastatic burden was significantly lower in treated mice (*p* = 0.0153, t test). **(E-F)** Tumor burden area (lung: *p* = 0.0315 and liver: *p* = 0.0040, both Mann-Whitney test) and number of tumors per mm^2^ (lung: *p* = 0.0024, t test and liver: *p* = 0.0400, Mann-Whitney test) were significantly lower in treated mice. Metastatic burden quantification of lung and liver tissue immunolabeled with human mitochondria antibody for overall tumor area (E) and number of tumors per mm^2^ (F) for individual mice.

## Discussion

Osteosarcoma remains a major clinical challenge largely due to its propensity to metastasize. While MYC has been suggested to be associated with aggressive disease (56), there are currently no known mechanisms of metastatic spread in osteosarcoma. The paucity of well-characterized metastasis models for this disease limits the ability to identify mechanisms of metastasis as well as the development of therapies specifically directed at preventing or treating metastatic spread. To address this need, we describe the development and characterization of a large panel of PDXs and PDX-derived cell lines which can be used to study osteosarcoma metastasis. We observed that the PDXs and PDX-derived cell lines largely share the genomic features of their original patient tumors and exhibit genomic complexity that mirrors what is observed in the human disease and is distinct from commercially available cell lines. Similarly, transcriptome clustering indicates that the PDXs and PDX-derived cell lines cluster with their tumor of origin, indicating close fidelity that is maintained over time. We also observed that PDX-derived cell lines cluster with the patient of origin samples rather than with other established commercially available cell lines, suggesting a distinct biology and perhaps a closer similarity to the human disease. Overall, the genomic and transcriptomic studies we performed strongly support the significant added value of developing new PDXs and PDX-derived cell lines for rare cancers rather than relying completely on the often-limited number of highly passaged immortal cell lines currently widely available.

We characterized the *in vivo* metastatic characteristics of eight PDX-derived cell lines and demonstrated that they have a wide phenotypic range with regard to primary OT growth and metastatic progression. The PDX-derived cell lines are heterogenous with respect to their tumorigenicity as well as their metastatic propensity and tropism. We anticipate that this will be a particularly useful feature of this resource as it will facilitate the selection of cell lines for the investigation of specific aspects of the osteosarcoma metastatic cascade. Three PDX-derived cell lines (OS742, OS152, and OS525) were found to have a consistently high metastatic capacity whereas two cell lines were non-metastatic (OS457 and OS186), and another three had an intermediate metastatic phenotype (OS052, OS526, and OS384). Of the cell lines tested *in vivo*, OS457 and OS186, were the PDX-derived cell lines most divergent from their patient tumor, due to WGD (Supplemental Figure S2A). It is possible that this drift has deleterious effects on the *in vivo* growth of these cell lines. PDX-derived cell lines metastasized most frequently to the lung, which is also the most common metastatic site in patients. Multiple PDX-derived cell lines also metastasize to bone (OS052, OS152, OS525, and OS742), another important metastatic site in osteosarcoma patients and an essentially non-existent feature in other previously described models, with the exception of the naturally occurring canine model (37, 45). The two most aggressive cell lines (OS152 and OS742) form invasive OT xenografts that reach the amputation endpoint in < 6 weeks and metastasize to lung, liver, bone, and other sites less than 1 month post-amputation; these latency characteristics make this an appealing model for mechanistic studies and preclinical therapeutic development. The variation in metastasis predilection sites between osteosarcoma PDX-derived cell line models illustrates the ability of these models to recapitulate the spectrum of the human disease.

In prior work, we demonstrated that the CDK inhibitor dinaciclib leads to decreased subcutaneous xenograft growth for osteosarcoma PDX-derived cell lines that carry MYC amplification (11). Here, we assessed whether dinaciclib could be used to decrease the progression of metastasis in an OT amputation model. This proof-of-concept experiment, using one osteosarcoma PDX-derived cell line (OS152), demonstrated that this model can be used effectively to test the effects of targeting genomic features to inhibit metastasis.

The PDX-derived cell lines described here were derived from both primary and metastatic sites. We did not find a clear association between patient of origin metastatic disease status and the *in vivo* metastatic phenotype or xenograft growth of the PDX-derived cell lines. The PDX-derived cell line exhibiting the fastest xenograft growth (OS742) and the non-tumorigenic cell line (OS457) were both derived from primary tumors. Similarly, of the two most metastatic cell lines, one is primary in origin (OS742) and the other originates from a metastatic site (OS152). The established commercially available osteosarcoma cell lines evaluated here were all derived from primary tumors and were either highly tumorigenic and produced heavy metastatic burden, MG63.3 and SJSA, or were non-tumorigenic and produced minimal metastatic burden (MG63). For SJSA and MG63.3, especially in the IV model, the metastatic burden induced in a short timeframe, ~1 month, caused significant mouse morbidity in the mice and presented challenges with burden quantification.

In patients, conventional osteosarcoma most commonly originates centrally in the metaphyseal region. In our OT model, cells are implanted paratibially to avoid direct seeding into the bloodstream via the bone marrow. However, pre-amputation microCT and post-amputation histologic examination of the affected limbs confirmed that all cell capable of forming xenografts also invade through the cortex. Confirming central invasion of the osteosarcoma PDX-derived cell line xenografts is probably critical for more accurate modeling of the human disease. In PDX paratibial implantation amputation models, metastasis incidence is reported to be lower in the mouse model than in human patients, ranging from 25-50% between PDXs (36, 51). Using PDX-derived cell lines in the OT model we achieved metastatic rates that were more in line with patient outcome; in three PDX cell lines the metastatic rate was 100% and a fourth cell line reached 88% in the OT model. In summary, we describe here a panel of new osteosarcoma PDX-derived cell lines that we believe will be of wide use to the osteosarcoma research community.

## Methods

### Patient sample procurement

All samples were reviewed by a pathologist (FKH or SC) and the diagnosis was confirmed as osteosarcoma. Samples were received fresh and were grossly evaluated by a pathologist (FKH or SC) for viable tumor tissue. A representative sample was reserved for PDX implantation and the remainder of the sample flash-frozen for DNA/RNA extraction.

### PDX and cell line generation

For PDX generation, 1 mm^3^ fragments of fresh or frozen-thawed patient tumor were dipped in Matrigel (Corning) and implanted under the renal capsule of NSG mice. Mice were monitored for tumor growth by abdominal palpation for up to 1 year post-implantation. Successfully engrafted tumors were allowed to reach 1-2 cm^3^ before the mouse was euthanized and tumor harvested. Tumors were digested in a collagenase buffer and filtered through a 70 μm filter. For PDX passaging, cells were implanted SQ in the flank of NSG mice (5 x 10^5^ cells) in 30 μL of MEM alpha and 20 μl Matrigel. PDX-derived cell lines were generated as described previously (11). Mycoplasma testing and STR analysis (Supplemental Table S3) were performed on cell lines prior to *in vitro* and *in vivo* use.

### Sequencing

Flash-frozen patient tumors and PDX-derived cell lines were placed into a 1.5 mL tube on ice and disrupted with a plastic tissue homogenizer. PDX tumors were digested to a single cell suspension in Collagenase digest buffer and sorted for human cells using anti human HLA-ABC (Biolegend, 555555), DNA and RNA were extracted using an AllPrep Kit (Qiagen, 80204) with QIAshredder (Qiagen, 79654). RNA was quantified with a Nanodrop 2000c (Thermo Fisher) and RNA integrity analyzed on an Agilent 5200 Fragment Analyzer. RNA sequencing was performed as described previously (11).

#### RNAseq

##### Hierarchical clustering

Counts were normalized using the edgeR package as log CPM with prior count set to 2 (57). Combat was then used as a batch correction for the sample source (patient, PDX, or cell line) (58). The top 80^th^ percentile most variant genes (based on coefficient of variance) were used for hierarchical clustering using the Manhattan method for distance followed by the Ward.D2 agglomeration method.

##### Data acquisition

TARGET and CCLE transcript per million (TPM) data were downloaded from the UCSC Treehouse Public Data repository (https://treehousegenomics.soe.ucsc.edu/public-data/). For the TARGET dataset, we used the Tumor Compendium v10 Public PolyA TPM expression and clinical datasets released in July 2019, and we filtered for osteosarcoma TARGET samples. For CCLE, we downloaded the Cell Line Compendium v2 TPM expression and clinical datasets released in December 2019 and filtered for osteosarcoma cell lines. From our dataset, patient tumor samples from which cell lines were also derived were excluded from the analysis.

The samples from TARGET, CCLE, and our dataset were processed using the UCSC TOIL RNA-seq pipeline (https://github.com/BD2KGenomics/toil-rnaseq) and the RSEM TPM values were used in the correlation analysis. For CCLE and our cell line data, the Toil pipeline was used in-house to generate RSEM gene-level Hugo counts for differential gene expression analysis (59).

##### Correlation analysis

We performed the correlation analysis as described in Yu et al. (60). Briefly, we selected the 5,000 most variable protein-coding genes by interquartile range (IQR) across the osteosarcoma patient samples. We then used these genes to calculate the Spearman’s correlation coefficient for each patient sample (TARGET and Sweet-Cordero lab) compared to each cell line (CCLE and Sweet-Cordero lab).

Whole genome sequencing and allele-specific copy number analysis

Alignment of FASTQ files, somatic variant calling, and structural variant calling was performed as previously described (11). Copy number analysis and purity correction was also performed as previously described (11), with the difference of allele-specific copy number (ASCN) being calculated using the B-allele frequency based upon germline variant allele frequency. For samples without germline data available, population-level common SNPs were obtained from dbSNP and used as an alternative to patient-specific germline variants. Concordance of copy-number across the genome was calculated by tallying the fraction of the genome with the same integer copy-number value between related samples. Concordance was calculated separately for major and minor alleles.

##### Somatic variant calling and Circos plot generation

Single nucleotide variants (SNVs) and small-scale insertions/deletions (indels) were called using Mutect2 (GATK v3.8) (61, 62) and Strelka (v2.9.10) (63). To filter out low-confidence calls, only the variants having at least 4 supporting reads and called by both Mutect2 and Strelka were retained. In addition, a variant was considered a true positive when the variant allele frequency (VAF) was greater than 5%, the sequencing depth was ≥20, and there were ≥3 sequence reads supporting the variant call. Somatic structural variants were called from WGS data by combining the output of Manta (v1.6.0) (64), DELLY (v0.8.1) (65), GRIDSS (v2.9.4) (66)and SvABA (v.1.1.3) (67). Consensus calls were made by comparing the output of GRIDSS and the other three tools, with a maximum allowed distance of 100bp as measured pairwise between breakpoints. We used two approaches to estimate the copy number profile, ploidy, and purity of a tumor sample from matched normal and tumor WGS data. Approach one implements PURPLE (v3.0) which combines B-allele frequency (BAF) from AMBER (v3.3), read depth ratios from COBALT (v1.8), somatic variant and structural variant calls. Approach two implements alleleCount (v4.2.1) (https://github.com/cancerit/alleleCount) for allele counting and ascatNgs (68, 69) for estimating tumor purity, ploidy, and copy number profile. Manual inspections were taken when there was a discrepancy between the two approaches. Circos plots were generated using PURPLE (v3.0) to plot final somatic copy number calls, point mutation calls, and structural variation calls.

### Cell culture

Eight PDX-derived cell lines (Table 1) and three established commercially available cell lines were used for the *in vivo* studies. SJSA and MG63 were obtained from the American Type Culture Collection (ATCC). MG63.3 was a gift from Dr. Robbie Majzner (Stanford University). All cells were maintained in a 37°C 5% CO2 humidified incubator and cultured in standard DMEM media (Corning) supplemented with 10% bovine growth serum (Hyclone SH30541.03) and 1% penicillin-streptomycin-glutamine. Stable luciferase-expressing cell lines were generated using lentiviral transduction (Addgene plasmid # 39196) followed by sorting for GFP using a Sony SH800 FACS Cell Sorter.

For the cell proliferation assay, cells were seeded in a 96-well plate and allowed to attach overnight. Plates were transferred into an Incucyte^®^ Live-Cell Analysis System (Sartorius, model S3), and whole well images were collected every 6 hours until wells reached 100% confluence. Incucyte software (Incucyte S3 2019A) was used to create confluence masks for each cell line. Confluence over time was used to identify a period of exponential cell growth and the linear portion of the curve was used to calculate doubling time in R statistical analysis software (70).

### *in vivo* studies

For each cell line, a minimum of 4 mice were used for establishment of model endpoint timing. Luciferase-expressing cell lines were used for all PDX-derived cell lines and SJSA. Both parental or luciferase-expressing cell lines were used for a subset of models (OS052, OS152, OS186, and OS526). Only the parental version was used for MG63 and MG63.3. For IV metastasis studies, 1 x 10^6^ cells were injected via the lateral tail vein in 100 μL of phosphate buffered saline (PBS); 1 x 10^5^ cells were used for SJSA. For luciferase-expressing cell lines, BLI was performed every 28 days to monitor metastatic progression using a PerkinElmer IVIS Spectrum Imaging System. Based on worsening clinical signs associated with metastases and/or BLI signal intensity an endpoint for each cell line model was determined.

To model spontaneous metastasis, paratibial OT implantation followed by limb amputation was employed. At implantation, the periosteum of the medial aspect of the mid-tibia was gently scored with a 30-gauge needle prior to injecting 1 x 10^6^ cells, 1 x 10^5^ for SJSA, in 10 μL of Matrigel. After xenograft detection, tumor growth was monitored with twice-weekly measurements until the limb diameter reached 1 cm, when the affected limb was amputated by coxofemoral disarticulation. Prior to amputation, limb and lung microCTs were performed using the Perkin Elmer Quantum GX2 microCT Imaging System; a pulmonary microCT was also performed prior to euthanasia. If no xenograft could be identified within six months of implantation, the cell line was deemed non-tumorigenic. Amputated limbs were demineralized in Immunocal (StatLab) for 24 hours prior to longitudinal bisection for histologic processing. An endpoint for each cell line was determined as it was for the IV model. Mice were euthanized 1-6 months, cell line dependent, post-injection for the IV model or post-amputation for the OT model.

For the dinaciclib study, luciferase-expressing OS152 cells were implanted orthotopically, as described above. BLI was performed initially 3-days post-amputation and dinaciclib treatment commenced the following day. All mice received treatment for 3 weeks at 20 mg/kg 5 times/week IP and BLI was performed every 7 days. Dinaciclib powder (MedChemExpress) was dissolved in DMSO to 50 mg/mL for stock solution. For working solution, the stock was diluted to 2.5 mg/mL with 20% beta cyclodextrin in PBS immediately before injection. Mice were weighed weekly. BLI signal was tracked over time using Living Image software (Perkin Elmer). BLI signal at endpoint was log-transformed and means were compared between the vehicle and control groups with a two-tailed t test, after testing for normality. Differences were considered significant when *p* < 0.05.

For all *in vivo* studies, lungs, liver, and tissue from additional metastatic sites were harvested for histological evaluation and metastatic burden analysis. Prior to collection, 5 mL of PBS was injected into the right ventricle of the heart and the lungs were inflated with 1 mL of formalin via the trachea. Tissues were fixed for 24-48 hours in formalin and then transferred to 70% ethanol prior to histologic processing.

### Immunohistochemistry

IHC was performed on formalin-fixed paraffin-embedded tissues mounted on glass slides. Standard heat-induced epitope retrieval using sodium citrate buffer (pH 6.0) was performed prior to H2O2 quenching of endogenous peroxidase activity. Blocking and antibody dilutions were made in 5% normal horse serum (Vector Labs), and anti-human mitochondria (1:1000; Abcam ab92824) was incubated overnight at 4°C in a humidified chamber. Slides were incubated in secondary horse anti-mouse antibody (Vector Labs) and avidin-biotin (Vectastain Elite ABC kit) for 30 minutes at room temperature. Signal was developed with DAB peroxide substrate (Abcam ab64238).

### Slide scanning and image analysis

Histology slides were scanned by the UCSF Histology & Biomarker Core at 10x using the Zeiss Axio Scan.Z1 Digital Slide Scanner. Scanned slides were imported into QuPath Quantitative Pathology & Bioimage Analysis software. Tissue was outlined using simple tissue detection prior to manual correction to exclude shadows, debris, and other tissue regions not meant for analysis. Training images were created from multiple regions across a selection of slides and were used to train the pixel classifier to identify DAB positive tissue. The pixel classifier was used to create tumor annotations across all slides with a minimum object size of 250 μm^2^. All annotations were checked manually for accuracy prior to quantification of tumor burden.

### Statistics

Metastatic burden was compared between the treatment and control groups with a two-tailed t-test for normally distributed data and the Mann-Whitney test for non-normally distributed data. Differences were considered significant when *p* < 0.05.

### Study approval

Written informed consent was collected from all patients or parents in the case of patients less than 18 years old. Using established guidelines (Belmont Report), institutional review board approval was granted from each participating institution (71). All procedures involving mice were conducted in accordance with protocols approved by the Stanford or UCSF IACUCs.

## Author contributions

ASC, CRS, ALK and LCS, conceived and designed the study. CRS, EPY, ALK, and KM conducted the experiments and collected data. ENV analyzed data. CRS and ASC analyzed and interpreted the data. CL, KY, MS, MRB, and AGL provided support for the computational analysis. SGL, HYL, AS, IB, MP, and ATS acquired patient consent and procured samples. LCS, ALK, and PD generated the PDXs. FKH and SJC confirmed the diagnosis of osteosarcoma in patient samples. RSA, DGM, MZ, and RW acquired the patient samples surgically. MS assisted with gene expression analysis. ASC and CRS wrote the manuscript.

## Acknowledgments

We thank the member of the UCSF Helen Diller Family Comprehensive Cancer Center (HDFCCC) Preclinical therapeutics core and animal core at UCSF for their useful advice throughout the development of this progress. Sequencing was performed at the UCSF Center for Advanced Technology (CAT), supported by UCSF PBBR, RRP IMIA, and National institutes of Health (NIH) 1S10OD028511-01 grants. Cell sorting was performed at the UCSF Laboratory for Cell Analysis (LCA), supported by the National Cancer Institute of the NIH under Award Number P30CA082103. The content is solely the responsibility of the authors and does not necessarily represent the official views of the NIH. CRS was supported by a Postdoctoral Research Fellow Grant from the Rally Foundation for Childhood Cancer Research. EASC was supported by the NIH (1R01CA243555) and by the Battle Osteosarcoma grant from St. Baldrick’s Foundation. Dr. Hani Goodarzi gifted the luciferase plasmid. Dr. Robbie Majzner of Stanford University gifted cell line MG63.3. Sara Pyle generated the graphical abstract.

