## Supplemental figures and tables for "Development and characterization of new patient-derived xenograft (PDX) models of osteosarcoma with distinct metastatic capacities"

**Supplemental Figure S1. Osteosarcoma patient-derived xenograft (PDX) histology after multiple passages.** The histologic features of osteosarcoma PDXs are maintained over multiple passages; OS106 maintains a fibroblastic phenotype (top panel), while OS525 maintains an osteoblastic phenotype (bottom panel). The histologic phenotype of the OS525 patient tumor was similar to that of multiple passages of its PDX and of the PDX-derived cell line xenograft (bottom panel). Hematoxylin and eosin.

**Supplemental Figure S2. Patient-derived xenograft (PDX)-derived cell line copy number comparisons to patient tumor and PDX. (A)** Genome copy-number (CN) concordance between patient-derived cell lines and the primary patient tumor and associated PDX. Concordance reported as the fraction of the genome with the same integer copy-number. Concordance was calculated separately for the major and minor alleles. The Jaccard score is the percent of the genome that has the same CN state between the two samples (delta = 0). The greater the Jaccard score, the more consistent the samples. **(B)** Example comparison between a primary tumor and the associated cell line. Top panel shows the allele-specific copy number (ASCN) for the tumor. Bottom panel shows the ASCN for the cell line. Middle panels show the change in CN for the major and minor alleles. Major alleles are in purple, minor alleles are in orange.

**Supplemental Figure S3. Circos plots showing structural rearrangements, copy number changes, and mutations for longitudinal samples.** Single nucleotide variants (SNVs), small-scale insertions/deletions (indels), and somatic structural variants, were called from WGS data. Somatic copy number calls, point mutation calls, and structural variant calls were used to generate Circos plots to compare the patient sample with the matched patient-derived xenograft (PDX) samples. **(A)** Structural rearrangements were similar between patient and PDX for OS384. **(B)** For OS052 the structural rearrangements were similar between patient, PDX, and PDX-derived cell line.

**Supplemental Figure S4. Metastatic capacity of osteosarcoma patient-derived xenografts (PDXs), spontaneously during subcutaneous (SQ) passaging, and after intravenous (IV) injection. (A)** Clusters of large neoplastic cells were identified on hematoxylin and eosin-stained sections (left) and were confirmed to be human osteosarcoma PDX metastasis using immunohistochemistry targeting human mitochondria antibody (right). **(B)** For most of the PDXs (8/11), lung metastases were detected during SQ passaging while maintaining the PDXs. **(C)** To evaluate the IV metastatic capacity of PDXs, $1 \times 10^6$ cells were injected into the lateral tail vein of NSG mice. Mice were euthanized after 28 days, and the lungs were harvested for burden quantification; 3/5 PDX-derived cell lines metastasized after IV injection. Three of the five PDXs tested metastasized to the lungs.

**Supplemental Figure S5. Comparison of parental and luciferase-expressing osteosarcoma patient-derived xenograft (PDX)-derived cell lines *in vitro* and in an *in vivo* intravenous (IV) model.** For all graphs, within each cell line, mice implanted at the same time are represented by the same symbol. **(A)** *in vitro* doubling time was similar between parental and luciferase expressing cell lines. **(B-D)** Metastatic burden and tropism varied slightly between parental and luciferase-expressing cell lines in the IV model. Burden quantification of lung tissue immunolabeled with human mitochondria antibody for overall tumor area (B) and number of tumors per $mm^2$ (C) for individual mice. IV model metastatic burden quantification of liver tissue immunolabeled with human mitochondria antibody for overall tumor area (D) and number of tumors per $mm^2$ (E) for individual mice.

**Supplemental Figure S6. Bioluminescence imaging (BLI) of mice injected intravenously (IV) with osteosarcoma patient-derived xenograft (PDX)-derived cell lines representing whole body burden over time.** 1 x $10^6$ cells were injected into the lateral tail vein of 2-5-month-old NSG mice and BLI was performed monthly until endpoint. For all graphs, within each cell line, mice implanted at the same time are represented by the same symbol. **(A-H)** BLI signal typically increased with time for the metastatic cell lines. Monthly BLI signal for each individual mouse for each cell line. **(I-J)** Several cell lines metastasized to bone. (I) BLI signal (left) identifying a metastatic lesion in the skull of a mouse injected with OS525; histology (right) confirmed the metastatic lesion was in the brain. Hematoxylin and eosin. (J) BLI signal (left) demonstrating lung and liver metastasis and identifying a metastatic lesion in the right hindlimb of a mouse injected IV with OS525; histology (right) confirmed the lesion was in the medullary cavity and trabecular bone of the proximal tibia. Hematoxylin and eosin.

**Supplemental Figure S7. Osteosarcoma patient-derived xenograft (PDX)-derived cell lines metastasize to the liver after intravenous (IV) injection.** For all graphs, within each cell line, mice implanted at the same time are represented by the same symbol. **(A-B)** Three cell lines were metastatic to liver, but also had lung metastases. Metastatic burden quantification of liver tissue immunolabeled with human mitochondria antibody for overall tumor area (A) and number of tumors per $mm^2$ (B) for individual mice. **(C)** Tumor size and distribution in the liver parenchyma varied between cell lines. Microscopic images of liver immunolabeled with human mitochondria antibody demonstrating metastatic burden for two PDX-derived cell lines.

**Supplemental Figure S8. Osteosarcoma patient-derived xenograft (PDX)-derived cell line xenograft growth after orthotopic implantation.** 1 x $10^6$ cells were implanted paratibially into the right hindlimb of 2–5-month-old NSG mice. After OT implantation, limb diameter was measured bi-weekly until reaching 1 cm when limb microCT was performed, followed by amputation via coxo-femoral disarticulation. **(A-H)** For the tumorigenic cell lines, limb diameter increased over time. OT xenograft growth for individual mice measured bi-weekly for each PDX-derived cell line. **(I)** Only one cell line, OS457, was non-tumorigenic**.** OS457 bioluminescence signal six months after implantation; xenografts were not palpable, nor could they be identified histologically. **(J)** PDX-derived cell line xenograft histopathology was heterogeneous for all tumorigenic cell lines; xenografts formed by individual cell lines vary in matrix production (top panel). Most patient of origin tumors (bottom panel) have regions with a similar histologic phenotype to the OT xenografts formed by their matched PDX-derived cell lines. Hematoxylin and eosin.

**Supplemental Figure S9. Comparison of parental and luciferase-expressing osteosarcoma patient-derived xenograft (PDX)-derived cell lines in an *in vivo* orthotopic (OT) model. (A-D)** Limb diameter increased with size after implantation; time to amputation was typically similar between parental and luciferase-expressing cell lines. Biweekly xenograft growth measurements from palpation to amputation in an OT model comparing parental to luciferase-expressing cells for individual mice. **(E-H)** Metastatic burden and site tropism was similar between parental and non-luciferase expressing cell lines. **(E-H)** For all graphs, within each cell line, mice implanted at the same time are represented by the same symbol. (E-F) OT model metastatic burden quantification of lung tissue immunolabeled with human mitochondria antibody for overall tumor area (E) and number of tumors per $mm^2$ (F) for individual mice. (G-H) OT model metastatic burden quantification of liver tissue immunolabeled with human mitochondria antibody for overall tumor area (G) and number of tumors per $mm^2$ (H) for individual mice. Only cell lines with burden are represented.

**Supplemental Figure S10. Bioluminescence imaging (BLI) representing whole body metastatic burden of mice post-amputation in an osteosarcoma patient-derived xenograft (PDX)-derived cell line orthotopic (OT) amputation model.** Metastatic capacity was determined by implanting 1 x $10^6$ cells paratibially into the right hindlimb of 2–5-month-old NSG mice, allowing growth to 1 cm in diameter, amputating the limb, and monitoring metastatic burden development using monthly BLI and health monitoring to determine study endpoint. At endpoint, tissues were harvested for quantitative burden analysis. For all graphs, within each cell line, mice implanted at the same time are represented by the same symbol. **(A-G)** BLI signal increased with time in the metastatic cell lines. Monthly post-amputation BLI signal for each individual mouse implanted with each PDX-derived cell line. **(H-I)** Several cell lines metastasized to bone. (H) BLI signal (left) demonstrating some liver metastases and identifying a metastatic lesion in the spine of a mouse implanted with OS525; histology (right) confirmed the metastatic lesion was in the vertebrae and had invaded out into the spinal canal. Hematoxylin and eosin. (I) BLI signal (left, middle) demonstrating liver metastasis and identifying widely distributed bone lesions in the left hindlimb (red box), spine, and skull (grey boxes) of a mouse implanted with OS152; histology confirmed the lesions were in the mid-to-distal femur (red solid boxes and bottom right hashed boxes), parietal bone (yellow hashed box, top left), and nasal turbinates (top right hashed boxes).

**Supplemental Figure S11. Osteosarcoma patient-derived xenograft (PDX)-derived cell lines metastasize to the liver after orthotopic (OT) injection.** For all graphs, within each cell line, mice implanted at the same time are represented by the same symbol. **(A-B)** Three cell lines were metastatic to liver. Metastatic burden quantification of liver tissue immunolabeled with human mitochondria antibody for overall tumor area (A) and number of tumors per $mm^2$ (B) for individual mice, only models that metastasized to liver are included. **(C)** Tumor size and distribution in the liver parenchyma varied between cell lines. Microscopic images of liver immunolabeled with human mitochondria antibody demonstrating metastatic burden for two PDX-derived cell lines.

**Supplemental Figure S12. Established commercially available osteosarcoma cell lines metastasize after intravenous (IV) injection.** IV metastatic capacity was determined by injecting 1 x $10^6$ (MG63 and MG63.3) or 1 x $10^5$ (luciferase-expressing SJSA) cells into the lateral tail vein of 2-5-month-old NSG mice. For all graphs, within each cell line, mice implanted at the same time are represented by the same symbol. **(A)** Total bioluminescence imaging (BLI) signal at endpoint representing whole body metastatic burden for SJSA. **(B)** BLI signal images at endpoint for SJSA demonstrating lung and liver metastasis. **(C-D)** Metastatic burden quantification of lung tissue immunolabeled with human mitochondria antibody for overall tumor area (C) and number of tumors per $mm^2$ (D) for individual mice. **(E)** Microscopic images of lung metastases immunolabeled with human mitochondria antibody demonstrating burden and growth pattern. **(F-G)** Metastatic burden quantification of liver tissue immunolabeled with human mitochondria antibody for overall tumor area (F) and number of tumors per $mm^2$ (G) for individual mice. **(H)** Microscopic images of liver metastases immunolabeled with human mitochondria antibody demonstrating burden.

**Supplemental Figure S13. Established commercially available osteosarcoma cell lines are tumorigenic and spontaneously metastasize in an orthotopic (OT) amputation model.** Metastatic capacity was determined by implanting 1 x $10^6$ (MG63 and MG63.3) or 1 x

$10^5$ (luciferase-expressing SJSA) cells paratibially into the right hindlimb of 2–5-month-old NSG mice, allowing growth to 1 cm in diameter, amputating the limb, and monitoring metastatic burden development using monthly bioluminescence imaging (BLI) and health monitoring to determine study endpoint. At endpoint, tissues were harvested for quantitative burden analysis. **(A-B)** All but one established cell line, MG63, formed xenografts that reached 1 cm in diameter. Bi-weekly xenograft diameter measurements for individual mice implanted with each osteosarcoma cell line. **(C-D)** Xenograft-bearing hindlimb microCTs with corresponding histopathology demonstrating medullary cavity and trabecular bone invasion and tumor expansion for the two tumorigenic established osteosarcoma cell lines. Hematoxylin and eosin. **(E)** Total BLI signal at endpoint for all SJSA mice, representing whole body metastatic burden. **(F)** BLI signal at endpoint for SJSA demonstrating lung and possibly liver metastasis. **(G-H)** Metastatic burden quantification of lung tissue immunolabeled with human mitochondria antibody for overall tumor area (G) and number of tumors per $mm^2$ (H) for individual mice. **(I)** Microscopic image of lung metastases immunolabeled with human mitochondria antibody demonstrating burden.

**Supplemental Table S1:** Established commercially available osteosarcoma cell line *in vivo* metastatic capacity in the literature

| **Cell Line**(derivation/original publication) | | | **Metastatic Capacity by Model**(publication) | | | | | | | |
|---|---|---|---|---|---|---|---|---|---|---|
| **Original** | **Altered/Passaged** | | **SQ/IM** | | **IV** | | **OT intra** | | **OT para** | |
| **MG63**(72, 73) | | | | No(74) | Yes(17) | No(12, 25) | Yes(27) | No(38) | | No(12) |
| | **MG63.2**(27) | | | | Yes(12) | | Yes(27) | | Yes(12) | |
| | **MG63.3**(12) | | | | Yes(12, 17, 18) | | | | Yes(12, 41) | |
| **U20S**(75) | | | | No(13, 74) | Yes(13) | No(25, 76) | | | | |
| | **U2OS/ MTX300**(76) | | | | | No(76) | Yes(77) | | | |
| **Saos2** | | | | No(74) | Yes(14, 19, 20) | No(12, 25) | Yes(20, 31, 32, 78–81) | No(38) | Yes(12) | |
| | **SaOS-LM1**(14) | | | | Yes(14) | | | | | |
| | **SaOS-LM2**(14) | | | | Yes(14) | No(15) | | | | |
| | **SaOS-LM3-6**(14) | | | | Yes(14, 15) | | | | | |
| | **SaOS-LM7**(15) | | | | Yes(12, 15) | | Yes(38, 40) | | Yes(12) | |
| **TE85**(82) | | | | | | No(12) | | No(38)* | | No(12) |
| | **HOS**(83) | | | No(74) | Yes(84) | No(12, 25) | Yes(80) | No(28)* | | No(12) |
| | | **MNNG/ HOS**(21) | | No(74, 85) | Yes(12) | | Yes(39) | | Yes(12) | |
| | | **KHOS/ R-970-5**(83) | Yes(85) | | | | | | | |
| | | **143B**(86) | Yes(74) | | Yes(12, 16) | | Yes(16, 28–30, 38, 39, 77, 87) | | Yes(12) | |
| | | **KRIB**(23) | | | Yes(12) | | Yes(34) | | Yes(12) | |
| **G292** | | | | No(74) | | No(25) | | No(38)* | | |
| **SJSA** | | | | No(74) | | | | | | |
| **OHS**(88) | | | | No(74) | | | | | | |

* = no xenograft formation, intra = intratibial implantation, para = paratibial implantation, Adapted and updated from(89)

**Supplemental Table S2:** *in vivo* metastatic characteristics of osteosarcoma PDX-derived cell lines, comparing luciferase-expressing to parental cells

| | | IV | | | | | | | | OT | | | | | | | | |
|---|---|---|---|---|---|---|---|---|---|---|---|---|---|---|---|---|---|---|
| Cell Line | | # Mice | Endpoint | Endpoint | Metastatic Site (%) | | | | | # Mice | Amp Time | Endpoint | Endpoint | Metastatic Sites (%) | | | | |
| PDX # | Version | [# Studies] | Decision | (avg d) | Any | Lung | Liver | Bone | Other | [# Studies] | (avg d) | Decision | (avg d) | Any | Lung | Liver | Bone | Other |
| OS152 | parental | 4 [1] | match luciferase | 48 | 100 | 100 | 100 | 0 | 0 | 7 [2] | 25 | match luciferase | 28 | 100 | 86 | 100 | 0 | 43 |
| | luciferase | 5 [2] | BLI, CS (abd mass) | 53 | 100 | 100 | 60 | 20 | 20 | 5 [1] | 39 | BLI | 29 | 100 | 100 | 100 | 40 | 0 |
| OS052 | parental | 5 [1] | match luciferase | 85 | 100 | 100 | 80 | 20 | 0 | 4 [1] | 30 | CS (abd mass) | 103 | 25 | 25 | 0 | 0 | 0 |
| | luciferase | 5 [2] | BLI | 85 | 100 | 100 | 0 | 20 | 20 | 9 [3] | 35 | max | 168 | 22 | 0 | 0 | 11 | 22 |
| OS526 | parental | 7 [2] | match luciferase | 140 | 100 | 100 | 0 | 0 | 0 | 6 [2] | 125 | match luciferase | 156 | 0 | 0 | 0 | 0 | 0 |
| | luciferase | 5 [2] | BLI | 140 | 100 | 100 | 0 | 0 | 0 | 5 [2] | 144 | max, CS (low BCS) | 159 | 0 | 0 | 0 | 0 | 0 |
| OS186 | parental | 5 [1] | alternate study | 112 | 0 | 0 | 0 | 0 | 0 | 5 [1] | 103 | C. bovis infection | 85 | 0 | 0 | 0 | 0 | 0 |
| | luciferase | 6 [2] | max | 160 | 0 | 0 | 0 | 0 | 0 | 7 [2] | 83 | max | 168 | 0 | 0 | 0 | 0 | 0 |

abd = abdominal, amp = amputation, avg = average, BLI = bioluminescence imagine, BCS = boody condition score, C. = Corynebacterium, CS = clinical signs, max = maximum

**Supplemental Table S3:** Short tandem repeat (STR) analysis of osteosarcoma PDX-derived cell lines

| Locus | OS152 | OS186 | OS457 | OS525 | OS526 | OS384 | OS742 | OS052 | OS774 | OS766 | OS833 |
|---|---|---|---|---|---|---|---|---|---|---|---|
| **AMEL** | X | X, Y | X, Y | X | X, Y | X | X | X | X | X | X,Y |
| **CSF1PO** | 10, 13 | 11, 12 | 10, 12 | 10 | 11, 13 | 11, 12 | 7, 11 | 11, 12 | 10, 13 | 10 | 11, 12 |
| **D13S17** | 11 | 8 | 10, 11 | 12 | 9 | 9 | 11, 14 | 13 | 11 | 12, 13 | 12 |
| **D16S539** | 11, 14 | 10, 13 | 10 | 12, 13 | 9, 12 | 12, 13 | 10 | 11 | 11 | 11, 14 | 11 |
| **D5S818** | 8, 12 | 10, 12 | 7, 13 | 11 | 11 | 11, 12 | 13 | 11 | 7, 12 | 13 | 10, 12 |
| **D7S820** | 10 | 10, 11 | 8, 14 | 10, 11 | 11 | 11, 12 | 8, 11 | 9, 11 | 10, 12 | 11 | 10, 11 |
| **TH01** | 9 | 7 | 6, 7 | 9.3 | 9 | 7 | 9 | 9.3 | 8 | 6, 9.3 | 7, 9.3 |
| **TPOX** | 8, 9 | 8, 11 | 8, 11 | 8, 11 | 8, 11 | 8, 11 | 8, 9 | 8 | 8 | 11 | 8 ,12 |
| **vWA** | 19, 20 | 16, 18, 19 | 16 | 16, 17 | 18 | 16, 18 | 17 | 17, 18 | 14, 16 | 17 | 15 |

**Supplemental Table S4:** Established osteosarcoma cell line *in vivo* intravenous (IV) study summary table

| | | MG63 | MG63.3 | SJSA |
|---|---|---|---|---|
| **INTRAVENOUS** | | | | |
| **# Mice [# Studies]** | | 8 [2] | 8 [2] | 5 [1] |
| **Endpoint (avg days)** | | 32 | 32 | 28 |
| **Endpoint Decision with CS, if Applicable** | | N/A | BLI, CS | BLI |
| | | N/A | respiratory signs | N/A |
| **Metastatic Sites (% )** | **Lung** | 100 | 100 | 100 |
| | **Bone** | 0 | 0 | 0 |
| | **Liver** | 0 | 0 | 100 |
| | **Other** | 0 | 0 | 0 |

abd = abdominal, avg = average, BCS = body condition score, BLI = bioluminescence imaging, CS = clinical signs

**Supplemental Table S5:** Established osteosarcoma cell line *in vivo* orthotopic (OT) study summary table

| | | MG63 | MG63.3 | SJSA |
|---|---|---|---|---|
| **ORTHOTOPIC** | | | | |
| **# Mice [# Studies]** | | 5 [1] | 4 [1] | 6 [1] |
| **Amp Time (avg days)** | | N/A | 36 | 31 |
| **Endpoint (avg days)** | | 247 | 29 | 28 |
| **Endpoint Decision with CS, if Applicable** | | N/A | BLI, CS | BLI |
| | | N/A | BCS | NA |
| **Metastatic Sites (%)** | **Lung** | 0 | 100 | 100* |
| | **Bone** | 0 | 0 | 0 |
| | **Liver** | 0 | 0 | 17* |
| | **Other** | 0 | 75 | 0 |

abd = abdominal, amp = amputation, avg = average, BCS = body condition score, BLI = bioluminescence imaging, CS = clinical signs, * = gross & BLI impression, tissues lost by courier

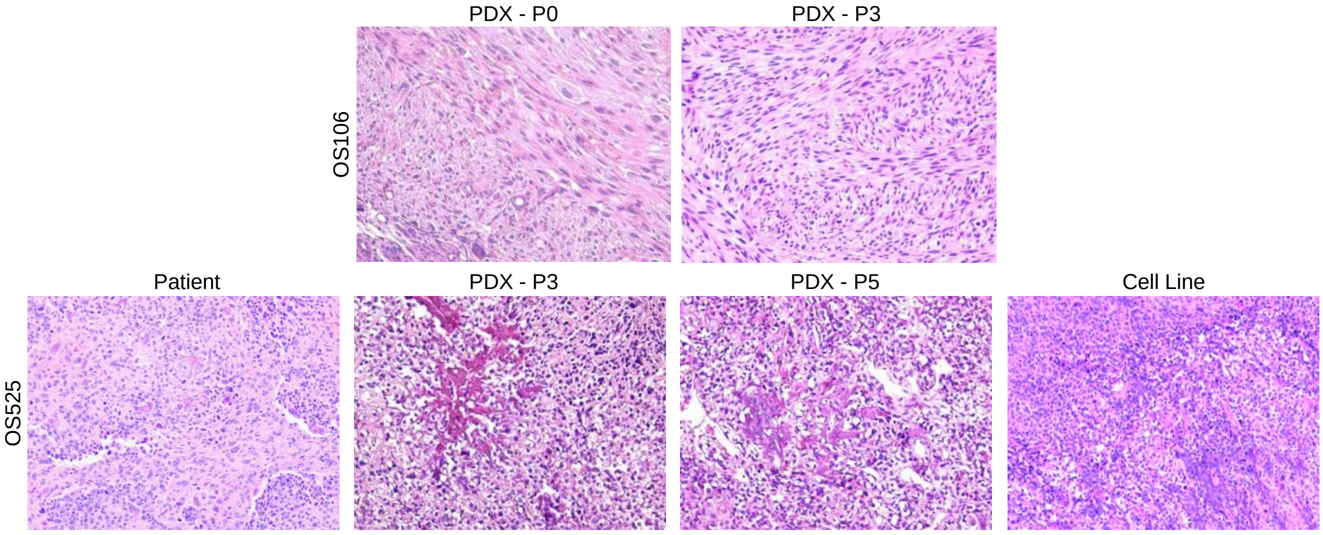


**S1**

A

B

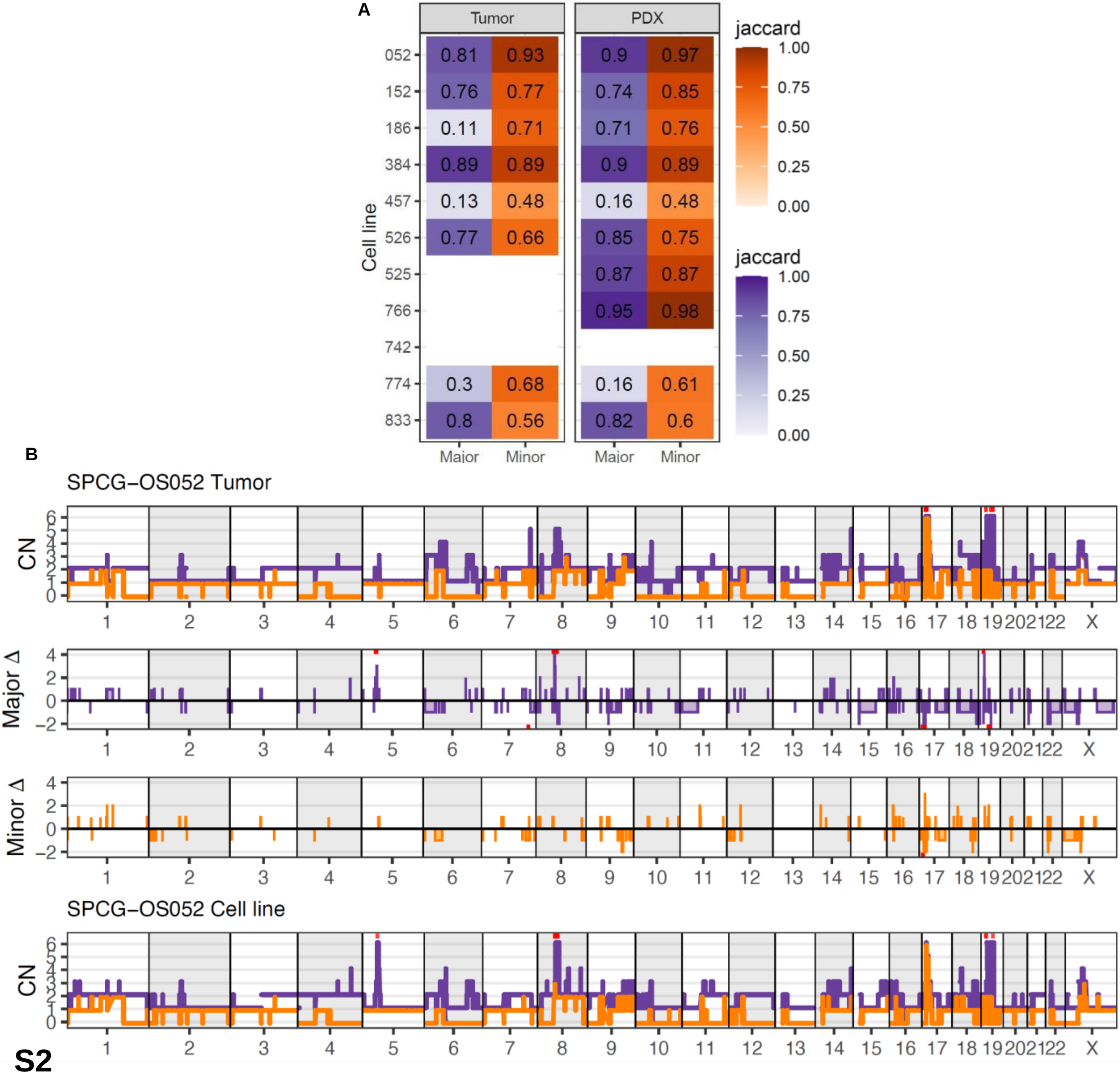
Tumor
PDX
jaccard
Cell line
Major
Minor
SPCG−OS052 Tumor
CN
Major Δ
Minor Δ
SPCG−OS052 Cell line

S2

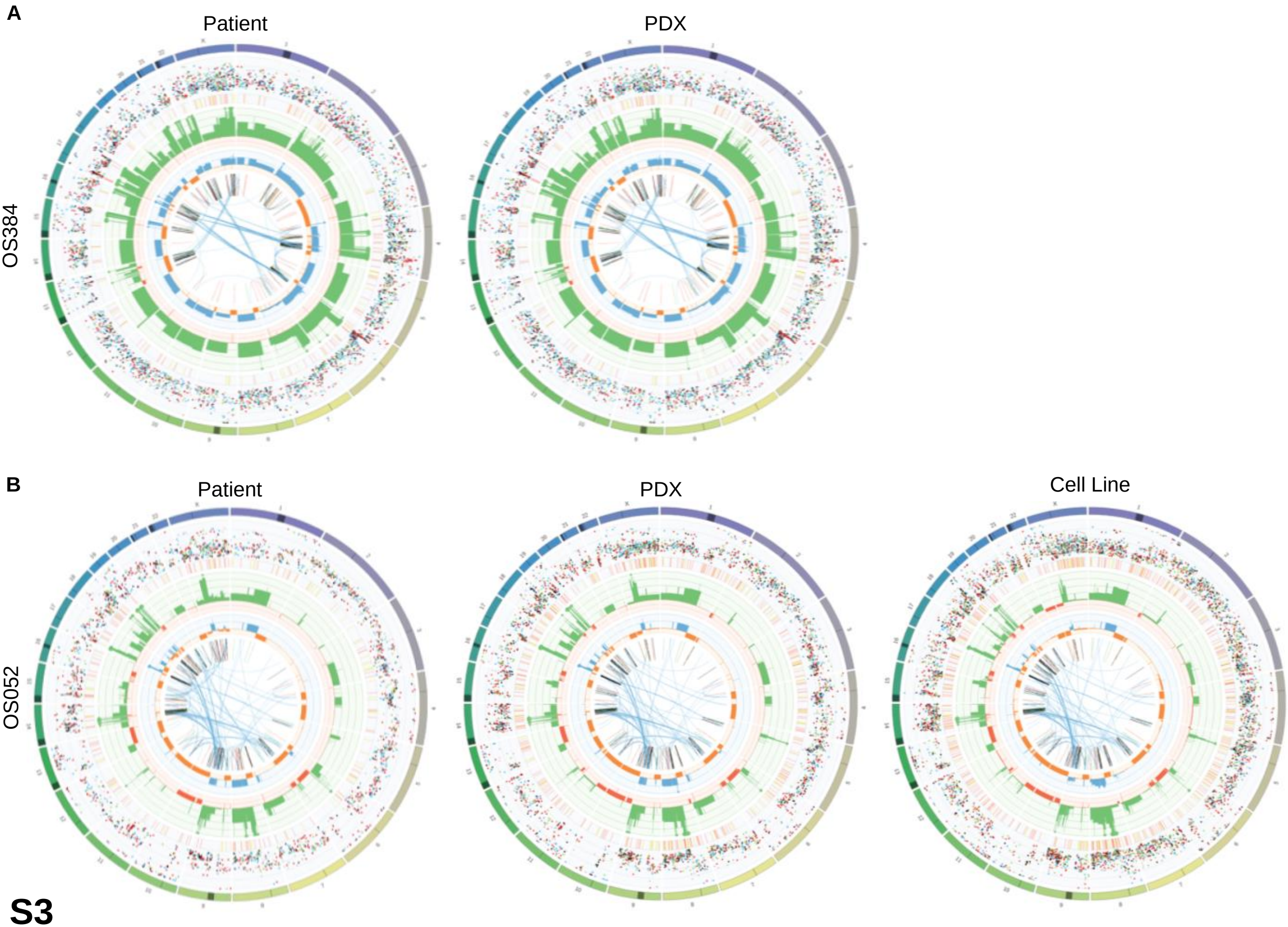


**S3**

A

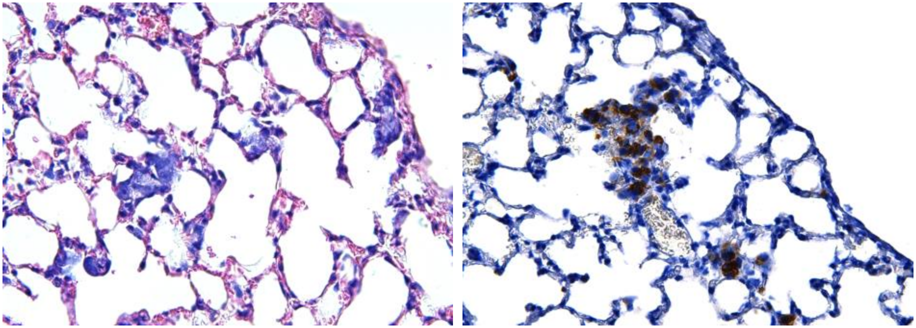

B

| PDX | Source | # Mice | % Metastasis |
|---|---|---|---|
| **OS052** | primary | 3 | 33 |
| **OS128** | primary | 1 | 0 |
| **OS566** | primary | 1 | 0 |
| **OS384** | primary | 3 | 33 |
| **OS457** | primary | 1 | 0 |
| **OS559** | primary | 1 | 100 |
| **OS106** | metastasis | 3 | 33 |
| **OS107** | metastasis | 7 | 71 |
| **OS152** | metastasis | 5 | 100 |
| **OS186** | metastasis | 11 | 64 |
| **OS337** | metastasis | 2 | 50 |

C

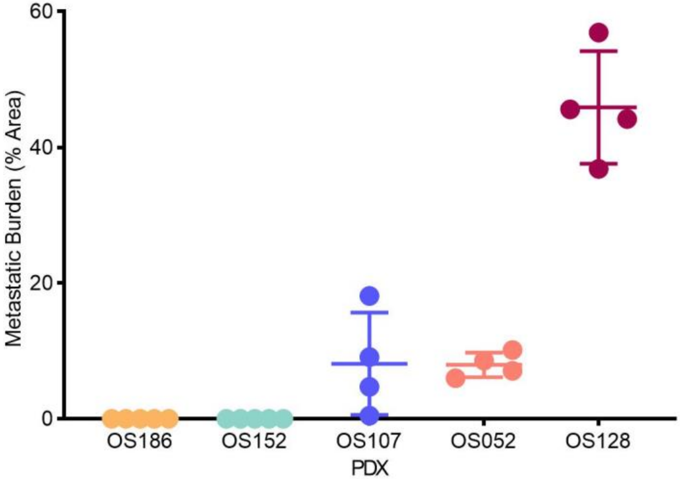


**S4**

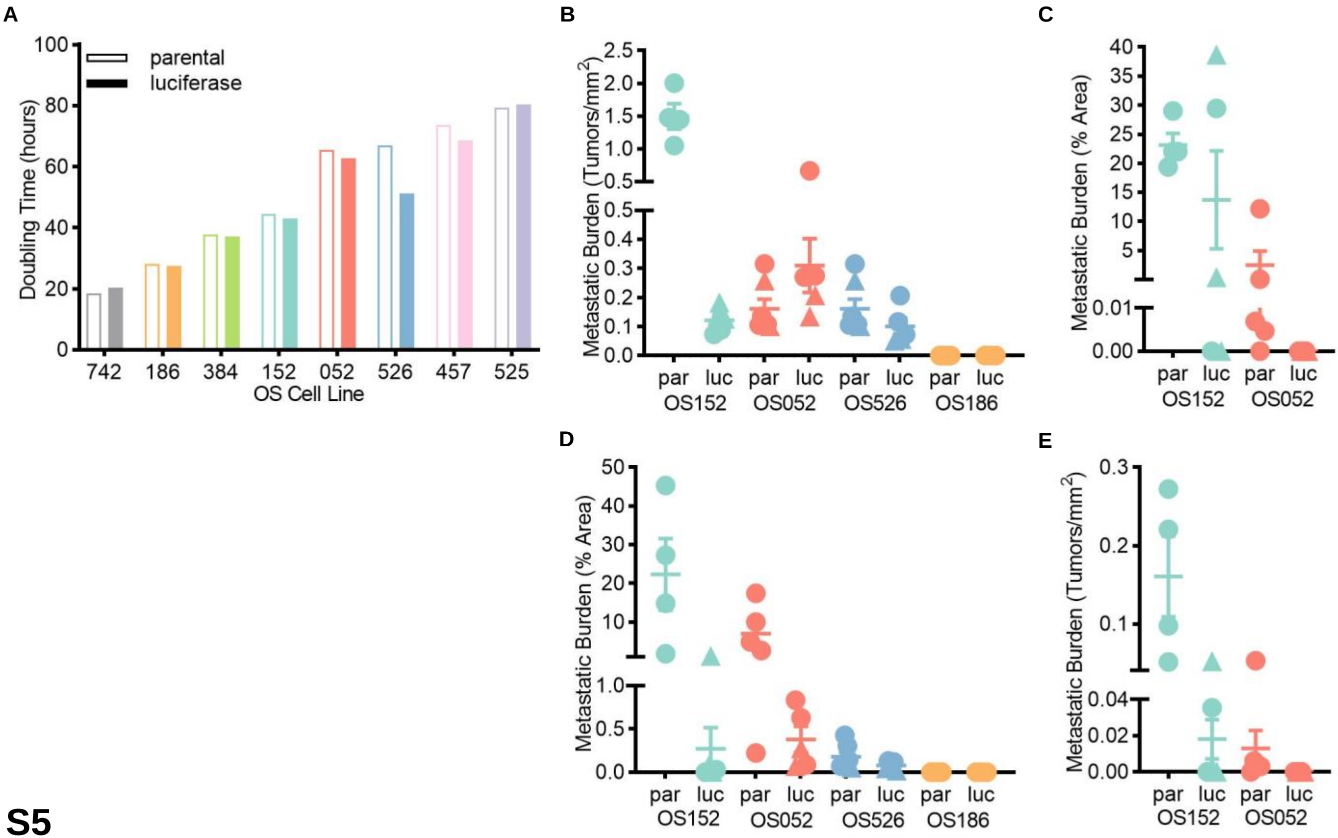

A
100
80
60
40
20
0
Doubling Time (hours)
parental
luciferase
742
186
384
152
052
526
457
525
OS Cell Line
B
Metastatic Burden (Tumors/mm2)
2.5
2.0
1.5
1.0
0.5
0.5
0.4
0.3
0.2
0.1
0.0
par
luc
par
luc
par
luc
par
luc
OS152
OS052
OS526
OS186
C
Metastatic Burden (% Area)
40
35
30
25
20
15
10
5
0.01
0.00
par
luc
par
luc
OS152
OS052
D
Metastatic Burden (% Area)
50
40
30
20
10
1.0
0.5
0.0
par
luc
par
luc
par
luc
par
luc
OS152
OS052
OS526
OS186
E
Metastatic Burden (Tumors/mm2)
0.3
0.2
0.1
0.04
0.02
0.00
par
luc
par
luc
OS152
OS052


**S5**

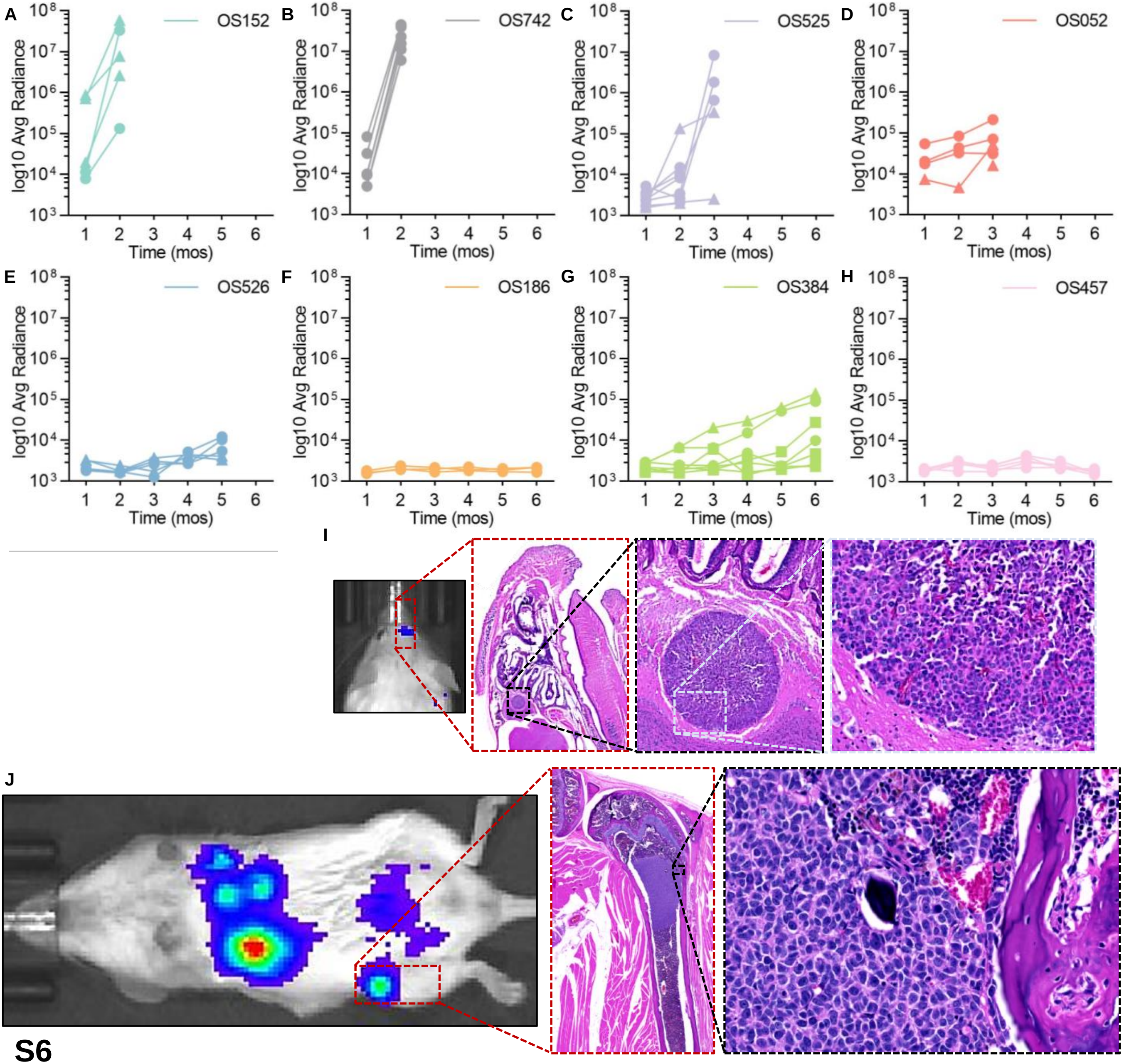
A
B
C
D
E
F
G
H
I
J
OS152
OS742
OS525
OS052
OS526
OS186
OS384
OS457
log10 Avg Radiance
Time (mos)
$10^3$
$10^4$
$10^5$
$10^6$
$10^7$
$10^8$
1
2
3
4
5
6

S6

A

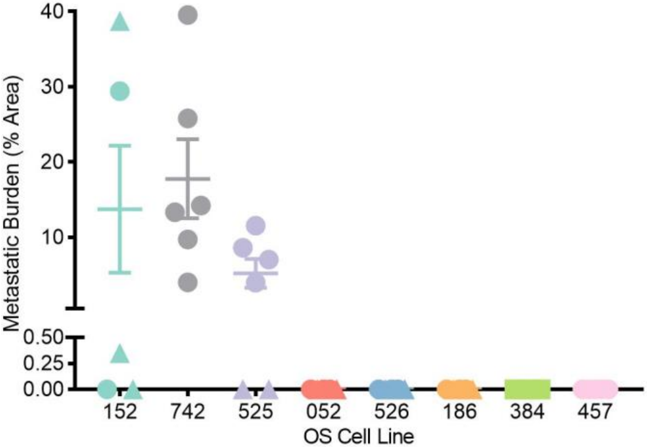


B

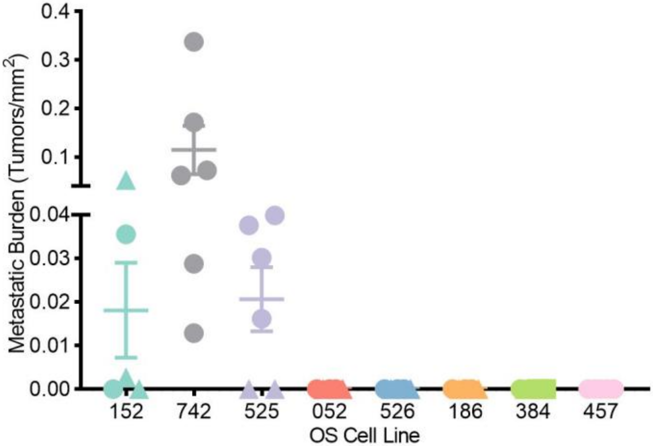


C

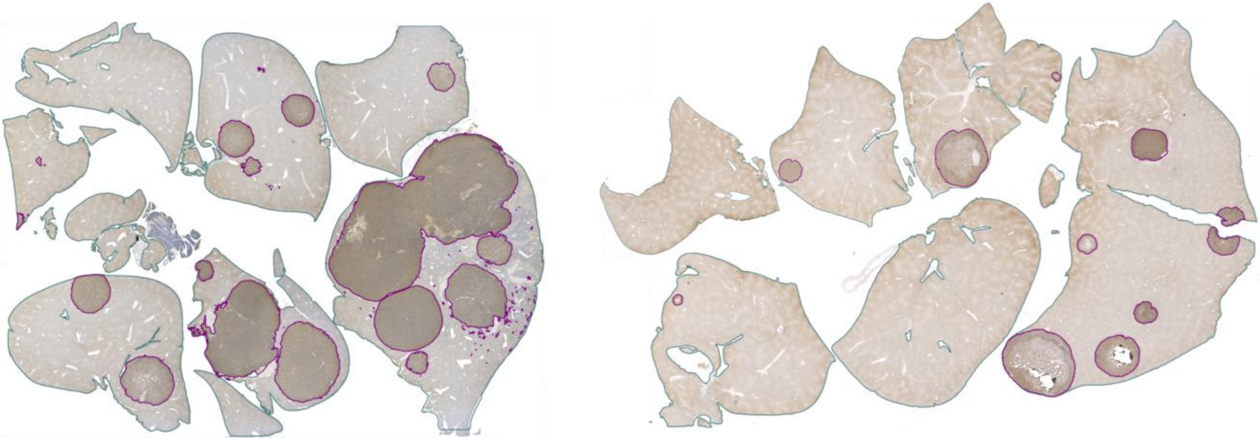

S7

OS152

OS525

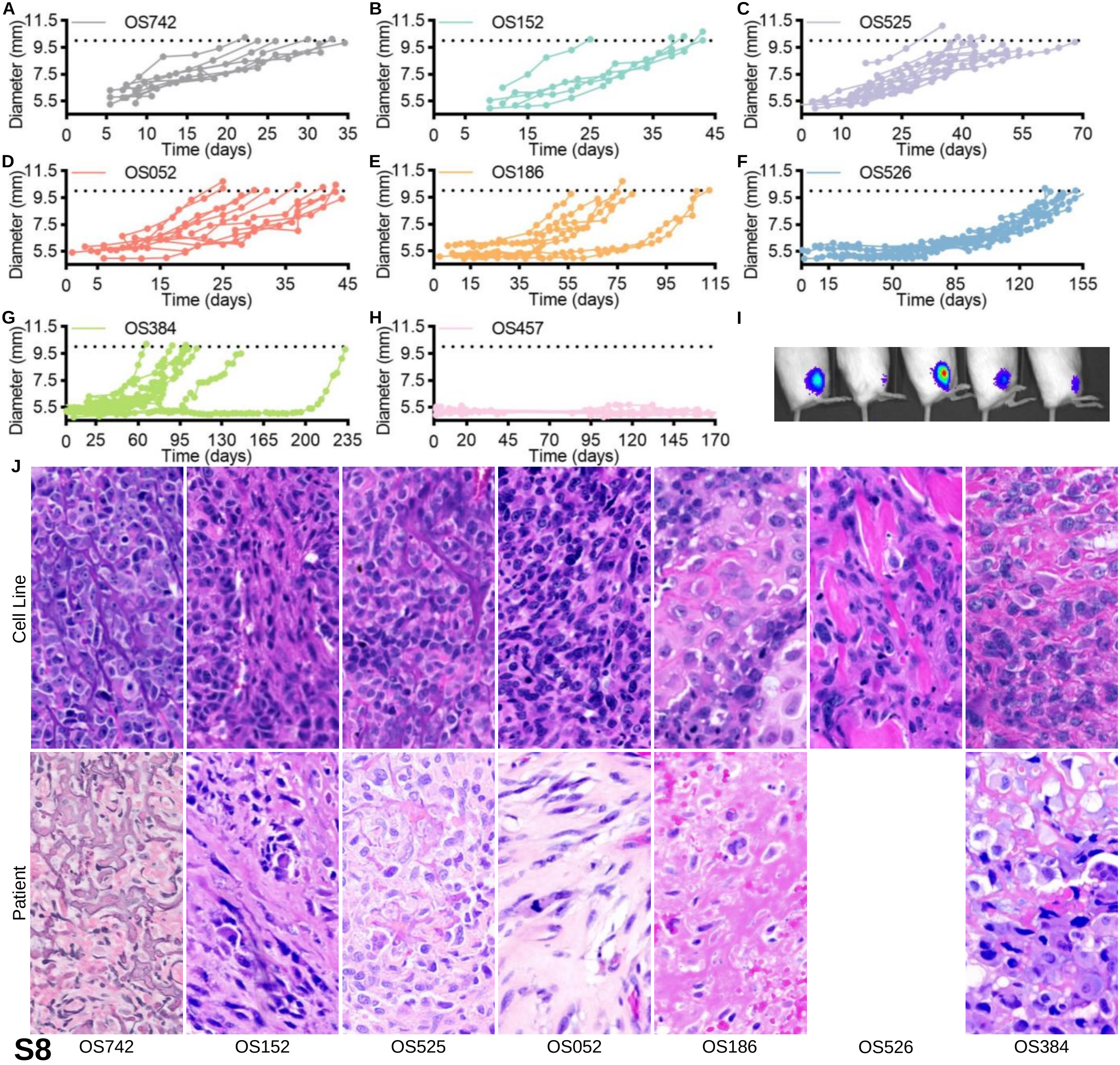
A
OS742
B
OS152
C
OS525
D
OS052
E
OS186
F
OS526
G
OS384
H
OS457
I
J
Diameter (mm)
Time (days)
Cell Line
Patient
OS742
OS152
OS525
OS052
OS186
OS526
OS384

**S8**

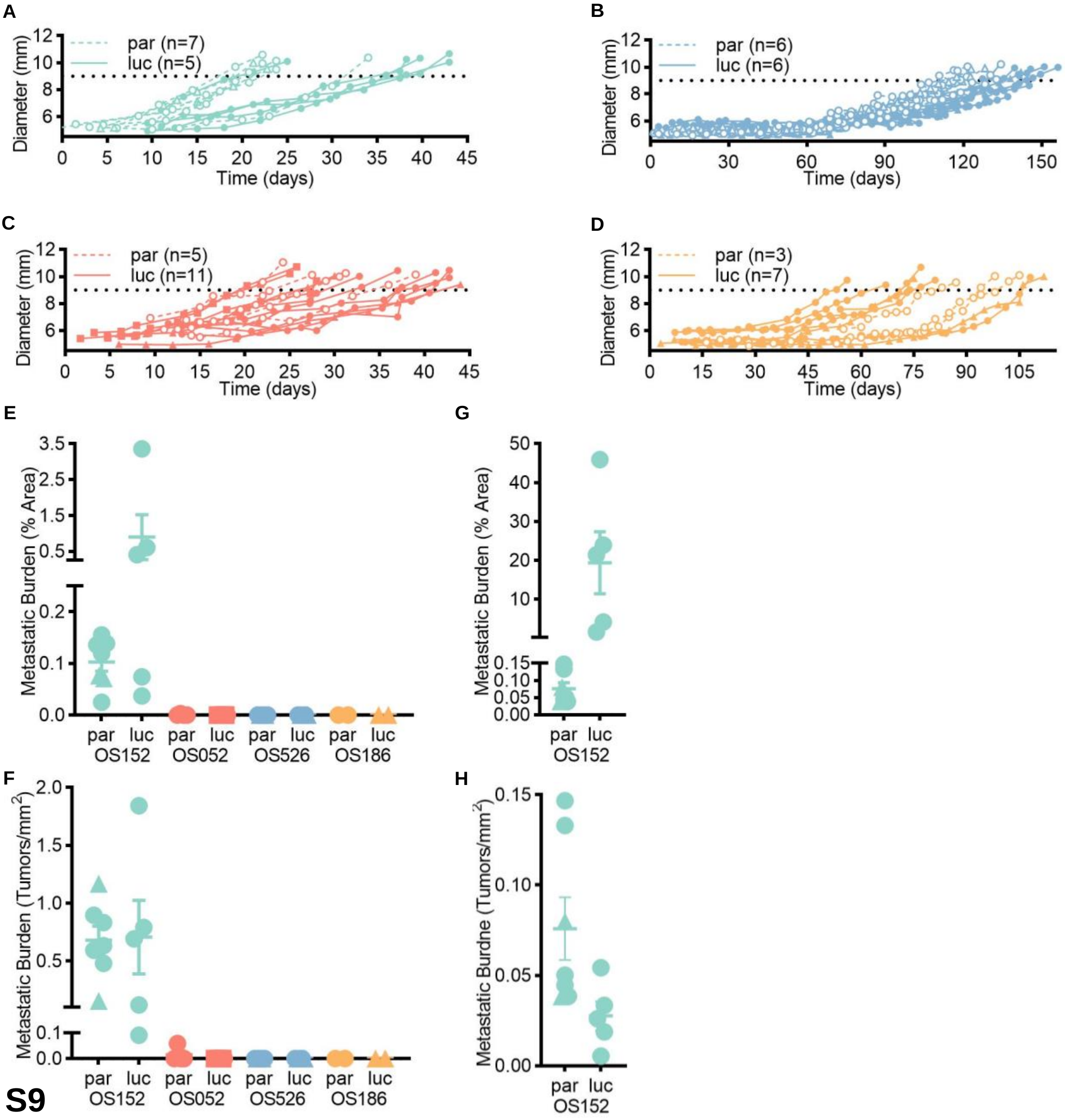
A
par (n=7)
luc (n=5)
Diameter (mm)
Time (days)
B
par (n=6)
luc (n=6)
Diameter (mm)
Time (days)
C
par (n=5)
luc (n=11)
Diameter (mm)
Time (days)
D
par (n=3)
luc (n=7)
Diameter (mm)
Time (days)
E
Metastatic Burden (% Area)
par luc par luc par luc par luc
OS152 OS052 OS526 OS186
G
Metastatic Burden (% Area)
par luc
OS152
F
Metastatic Burden (Tumors/mm2)
par luc par luc par luc par luc
OS152 OS052 OS526 OS186
H
Metastatic Burdne (Tumors/mm2)
par luc
OS152

S9

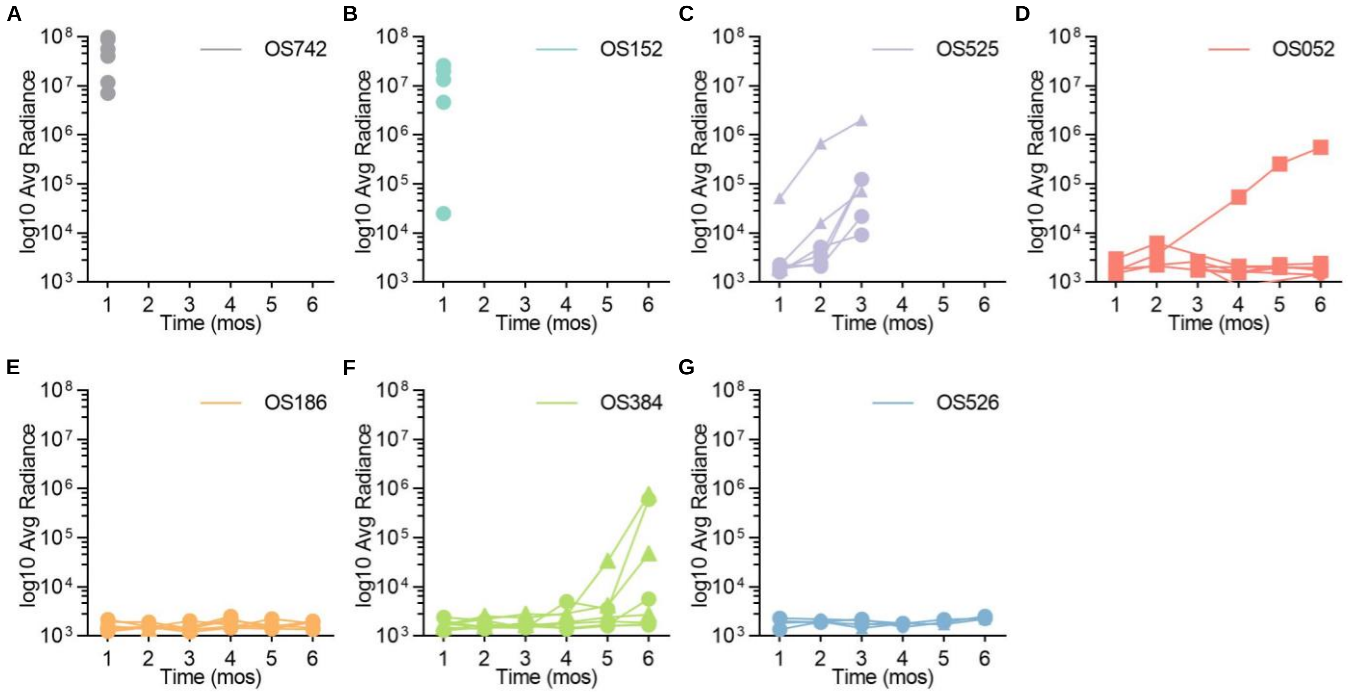
A
B
C
D
E
F
G
OS742
OS152
OS525
OS052
OS186
OS384
OS526
log10 Avg Radiance
Time (mos)
$10^3$
$10^4$
$10^5$
$10^6$
$10^7$
$10^8$
1
2
3
4
5
6

**S10**

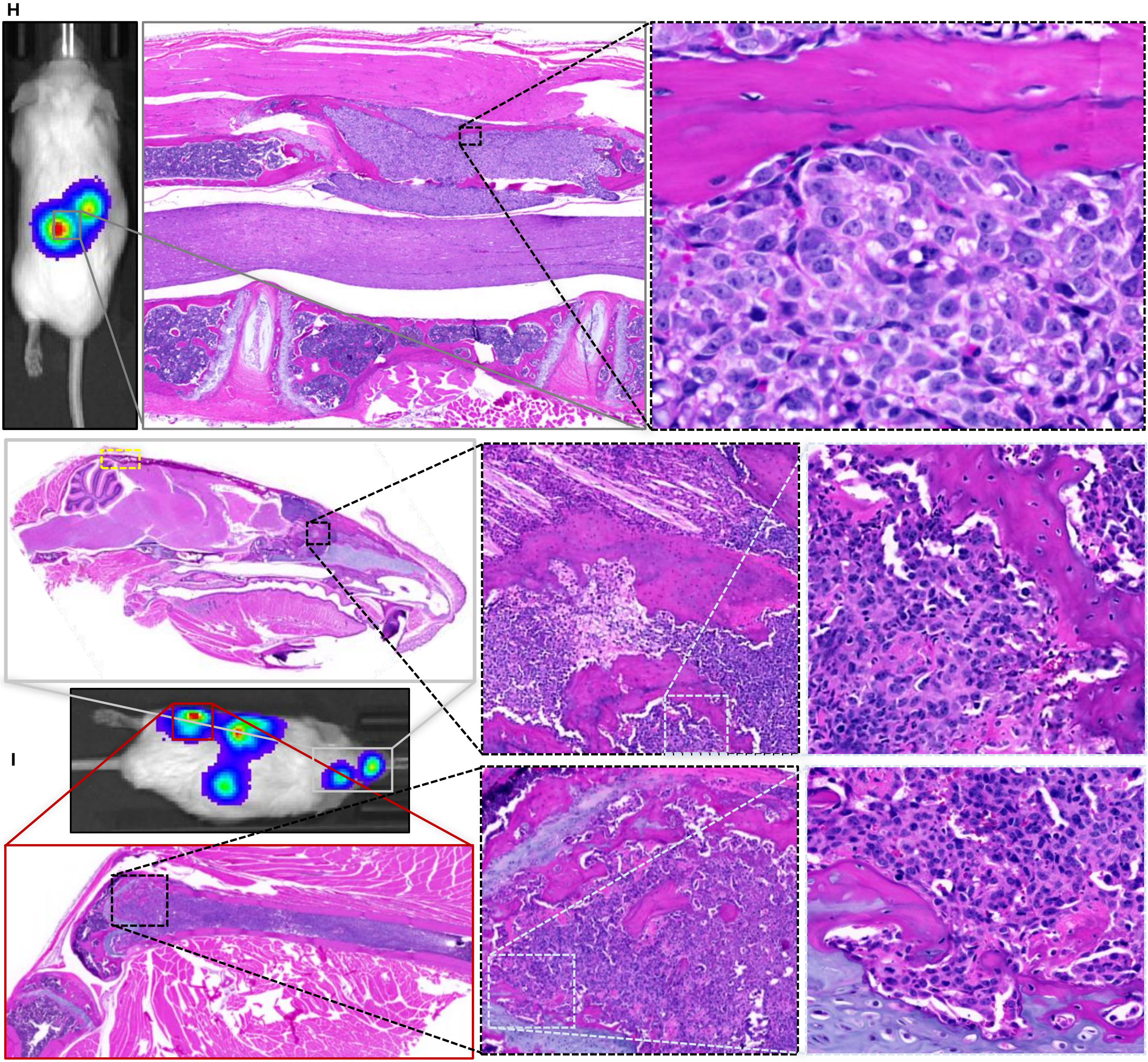


S10 continued

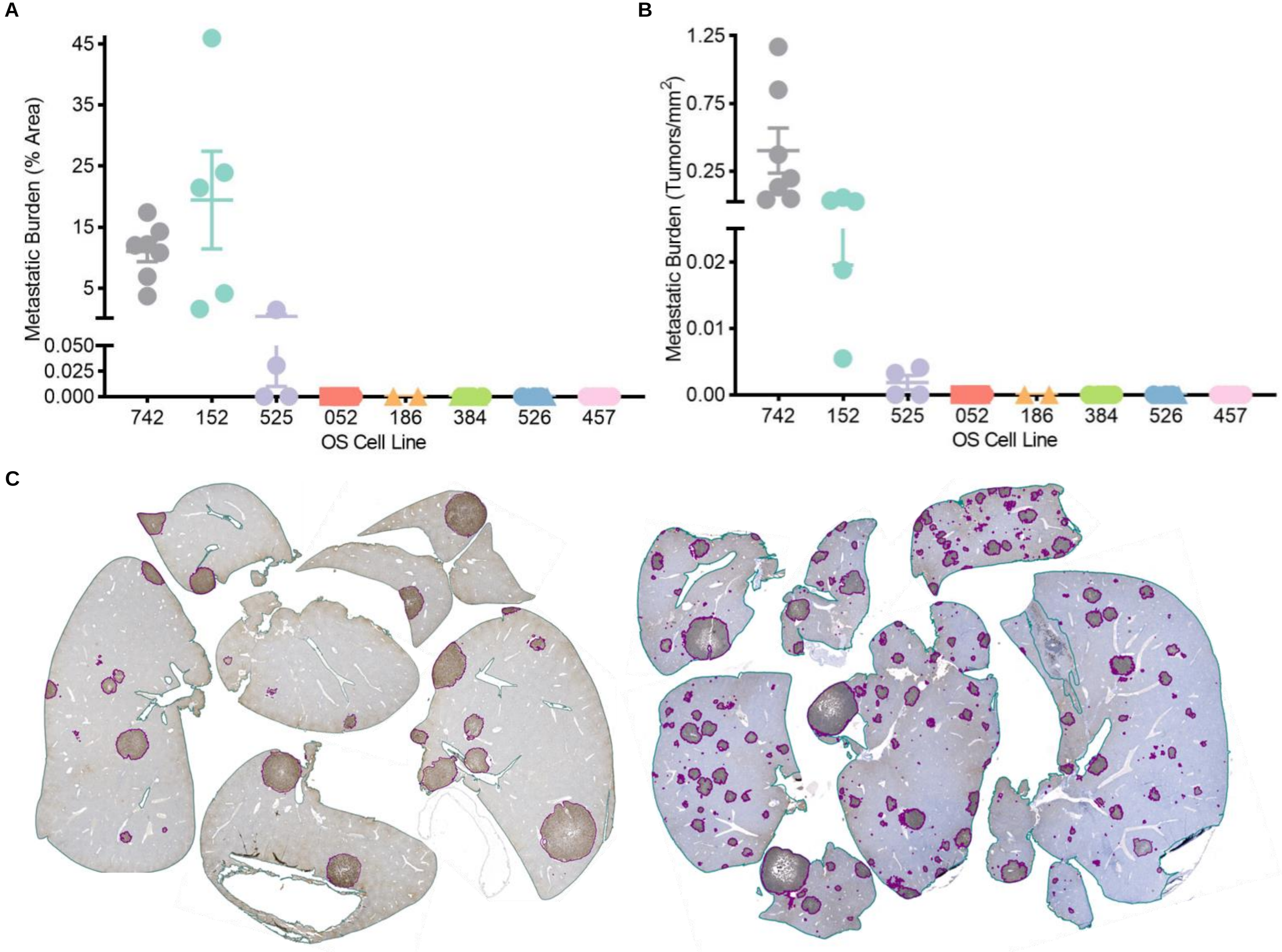


**S11**

OS152

OS742

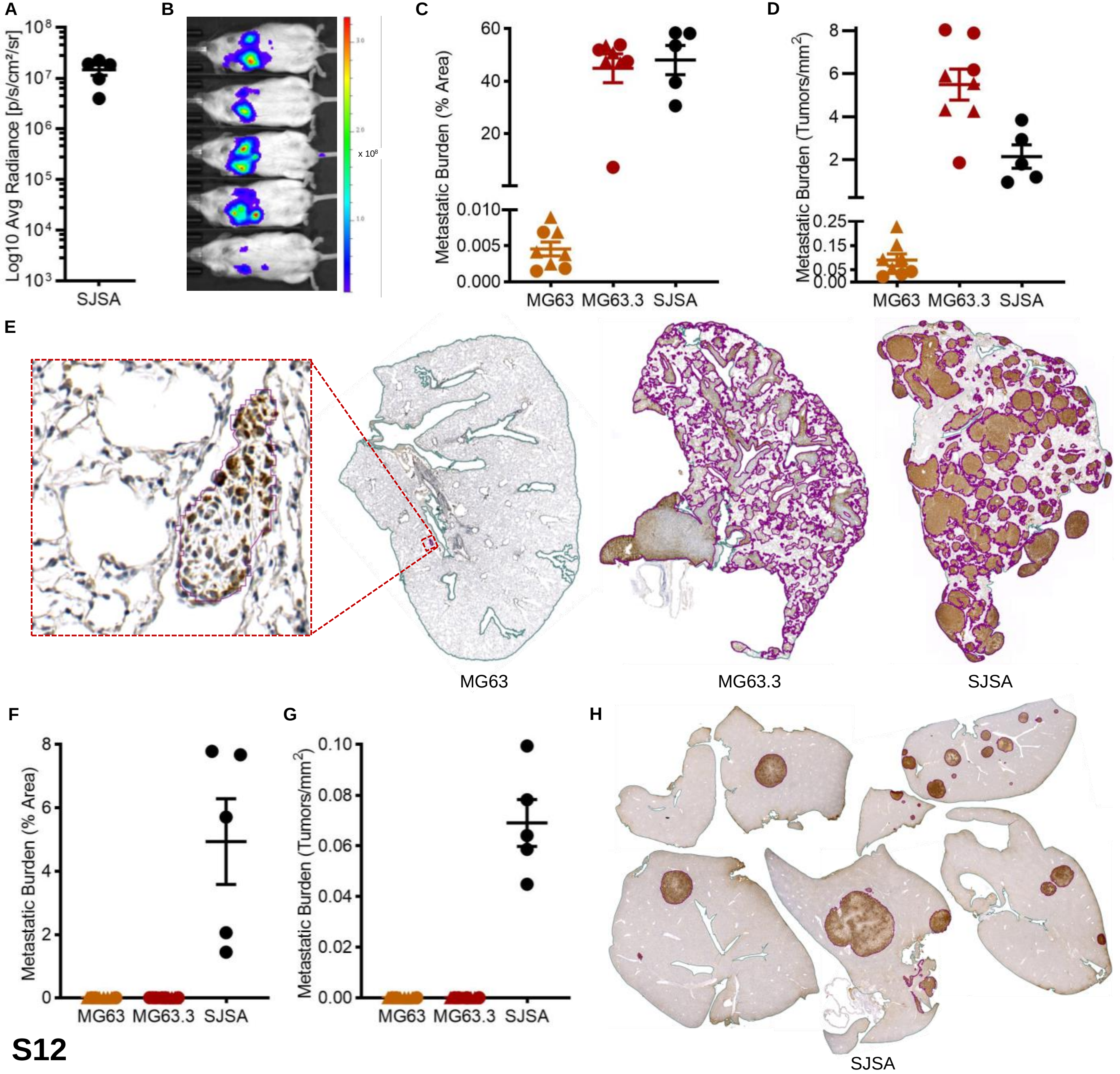

A
Log10 Avg Radiance [p/s/cm²/sr]
SJSA
B
x 10^8
C
Metastatic Burden (% Area)
MG63
MG63.3
SJSA
D
Metastatic Burden (Tumors/mm²)
MG63
MG63.3
SJSA
E
MG63
MG63.3
SJSA
F
Metastatic Burden (% Area)
MG63
MG63.3
SJSA
G
Metastatic Burden (Tumors/mm²)
MG63
MG63.3
SJSA
H
SJSA


**S12**

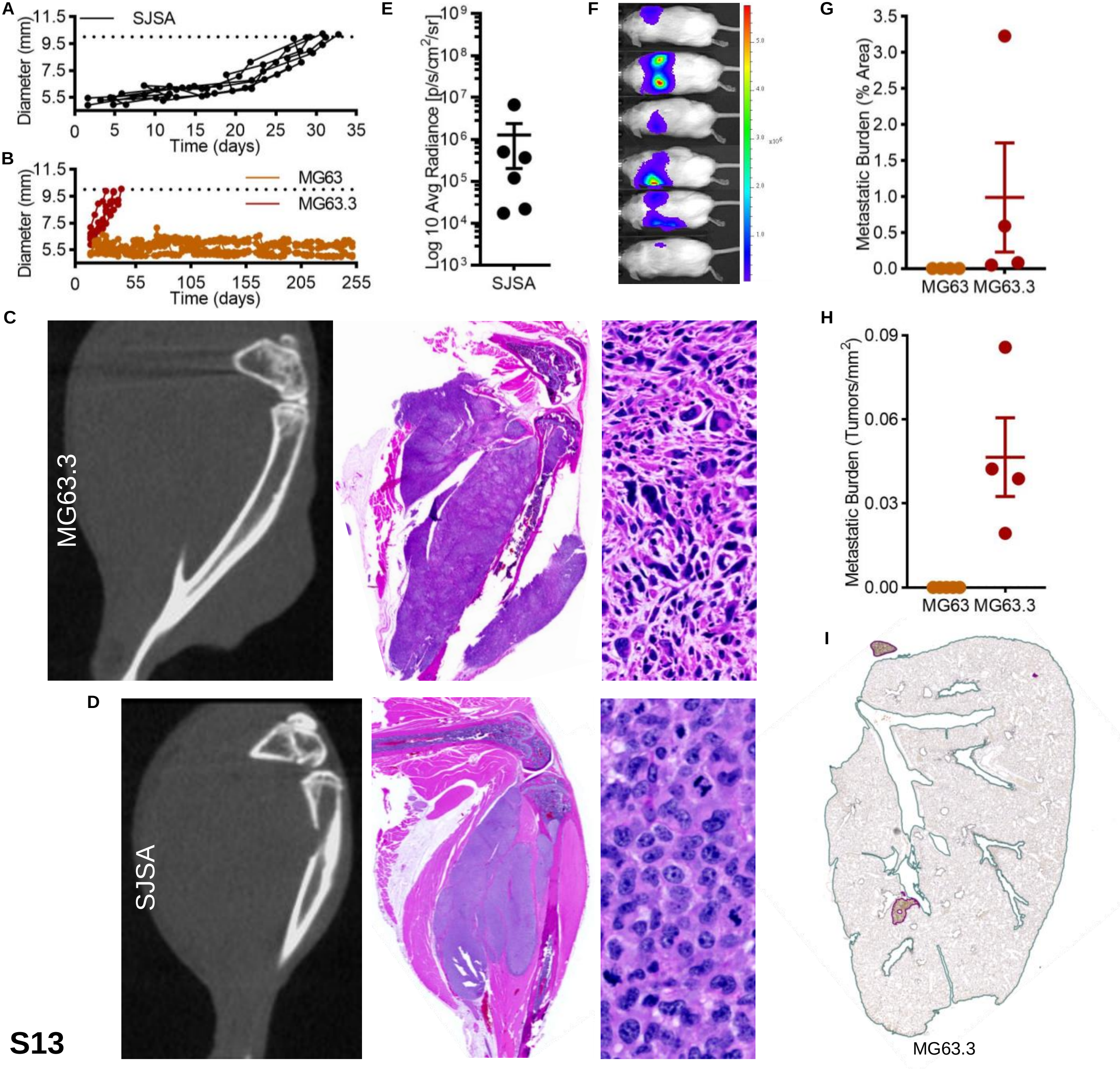

A
SJSA
Diameter (mm)
11.5
9.5
7.5
5.5
0
5
10
15
20
25
30
35
Time (days)
B
MG63
MG63.3
Diameter (mm)
11.5
9.5
7.5
5.5
0
55
105
155
205
255
Time (days)
E
Log 10 Avg Radiance [p/s/cm2/sr]
10^9
10^8
10^7
10^6
10^5
10^4
10^3
SJSA
F
5.0
4.0
3.0
x10^6
2.0
1.0
G
Metastatic Burden (% Area)
3.5
3.0
2.5
2.0
1.5
1.0
0.5
0.0
MG63
MG63.3
C
MG63.3
H
Metastatic Burden (Tumors/mm2)
0.09
0.06
0.03
0.00
MG63
MG63.3
D
SJSA
I
MG63.3

S13
